# A DNA-binding protein senses DNA superhelicity to switch between bridging and nucleoprotein filament formation

**DOI:** 10.64898/2026.05.23.727375

**Authors:** Sneha Shahu, Sreeja Baira, Sudhish Gupta, Mahipal Ganji

**Author notes:** Equal contribution (Sneha Shahu, Sreeja Baira).

## Abstract

The torsional state of DNA encodes regulatory information beyond sequence, yet how chromosome-associated proteins read this mechanical signal to switch between distinct functional outputs remains poorly understood. Using real-time single-molecule fluorescence imaging on topologically constrained DNA, we demonstrate that DNA superhelical polarity is the primary determinant of H-NS binding mode with direct consequences for chromosomal domain organization and gene silencing. On positively supercoiled DNA, H-NS assembles into nucleoprotein filaments at AT-rich loci, acting as a topological barrier that confines plectoneme diffusion. On negatively supercoiled DNA, H-NS switches to bridging mode, immobilizing plectonemes at AT-rich sites. Upon changes in DNA topology, H-NS dynamically switches between binding modes within seconds. An oligomerization-deficient mutant retains bridging but cannot form filaments, confirming the mechanistic distinction. On multi-AT-locus DNA, helicity-driven mode selection produces self-organized “chromosomal” architecture: some sites stochastically capture the plectoneme and bridge, while remaining sites assemble insulating filaments. These findings establish DNA superhelical polarity as the master switch governing H-NS binding mode and “chromosomal” domain organization, with broad implications for transcriptional regulation in bacteria.

## Main text

Bacterial genomes are dynamically reorganized to balance the expression of core housekeeping genes while simultaneously silencing foreign DNA acquired through horizontal gene transfer. Among the proteins that mediate this organization, the nucleoid-associated protein H-NS in *Escherichia coli* plays a central role by selectively binding AT-rich sequences typical to horizontally acquired pathogenicity islands^1,2^. In doing so, H-NS functions both as a global transcriptional regulator and as a local structural organizer of the nucleoid, shaping chromosomal architecture locally through its ability to compact and loop DNA^3-6^. H-NS achieves its dual functions through two mechanistically distinct binding modes. In the *stiffening mode*, H-NS polymerizes cooperatively along the minor groove of AT-rich DNA, forming a rigid nucleoprotein filament that likely plays a structural role in chromosome organization^7,8^. In the *bridging mode*, H-NS cross-links two DNA duplexes in trans, generating compacted loop structures that sterically occlude RNA polymerase and silence transcription at H-NS target genes^9,10^. Both modes have been characterized structurally and biochemically^11-13^, yet despite these efforts, the question of what physiological signal selects one mode over the other has remained unresolved. Because of this, our understanding of how H-NS coordinates transcriptional silencing and chromosome organization has remains limited.

Previous work proposed that magnesium ion concentration drives H-NS binding mode switching^12,13^, where low Mg^2+^ favors stiffening^8^ and high Mg^2+^ promotes bridging^9^. However, follow-up studies suggested that elevated Mg^2+^ levels simply reduce H-NS DNA-binding affinity, with apparent bridging arising indirectly from reduced protein occupancy^14,15^, indicating that divalent cations as an unlikely physiological switch. Early biochemical evidence showed that H-NS constrains negative supercoiling in plasmid DNA^16^, and in *Salmonella*, H-NS was dislodged from chromosomal binding sites upon transcriptional activation of upstream genes, with the effect propagating at a distance and suppressed by a gyrase binding site^17^ – pointing to a dynamic coupling between transcription-driven supercoiling and H-NS occupancy, consistent with findings that H-NS binding sites are enriched in negatively supercoiled chromosomal regions^18^. As a constitutively expressed nucleoid-associated protein, H-NS will inevitably encounter both positive and negative supercoiling states generated by transcription and topoisomerase activity^19,20^. Yet how H-NS responds to each supercoiling state has remained unexplored.

Here we address this gap using real-time single-molecule fluorescence imaging that enables simultaneous visualization of unlabeled H-NS and surface-tethered supercoiled DNA of defined sequence. By exploiting the fluorescent DNA intercalating dye SYTOX Orange (SxO) to specifically generate positively or negatively supercoiled DNA *in situ*^21,22^, we directly visualized H-NS binding dynamics on topologically defined substrates in real time. Our experiments reveal that DNA superhelical polarity governs H-NS binding mode. Neither ionic conditions, nor DNA sequence alone determine this switch. Instead, positive supercoiling drives filament formation, and negative supercoiling drives bridging. H-NS can sense and switch between modes within seconds responding to topology changes, and an oligomerization-deficient mutant H-NS_Y61DM64D_ confirms that the two modes are mechanistically distinct and can be uncoupled. On multi-AT-locus DNA, this helicity-sensing mechanism results in a self-organized topological domain architecture whose functional implications potentially extend to gene silencing, topological insulation, and stochastic gene regulation in bacterial populations.

## Results

### H-NS forms nucleoprotein filaments at AT-rich loci on relaxed DNA

To establish a label-free single-molecule readout of H-NS binding, we constructed a 21 kb DNA with a 3 kb, 65% AT-rich segment at its center (Template-1, Fig. 1a). Both ends were ligated with 500 bp biotinylated DNA handles for dual-end immobilization on PEG-passivated glass via biotin–streptavidin linkage^21^ (Fig. 1b-1d). DNA was visualized using the intercalating fluorophore SYTOX Orange (SxO), which maintains rapid exchange kinetics that minimize photodamage and binds DNA without sequence preference^22,23^. Relaxed DNA in the absence of H-NS displayed uniform fluorescence along its length (Fig. 1e-left, Movie S1, S5).

**Figure 1.**
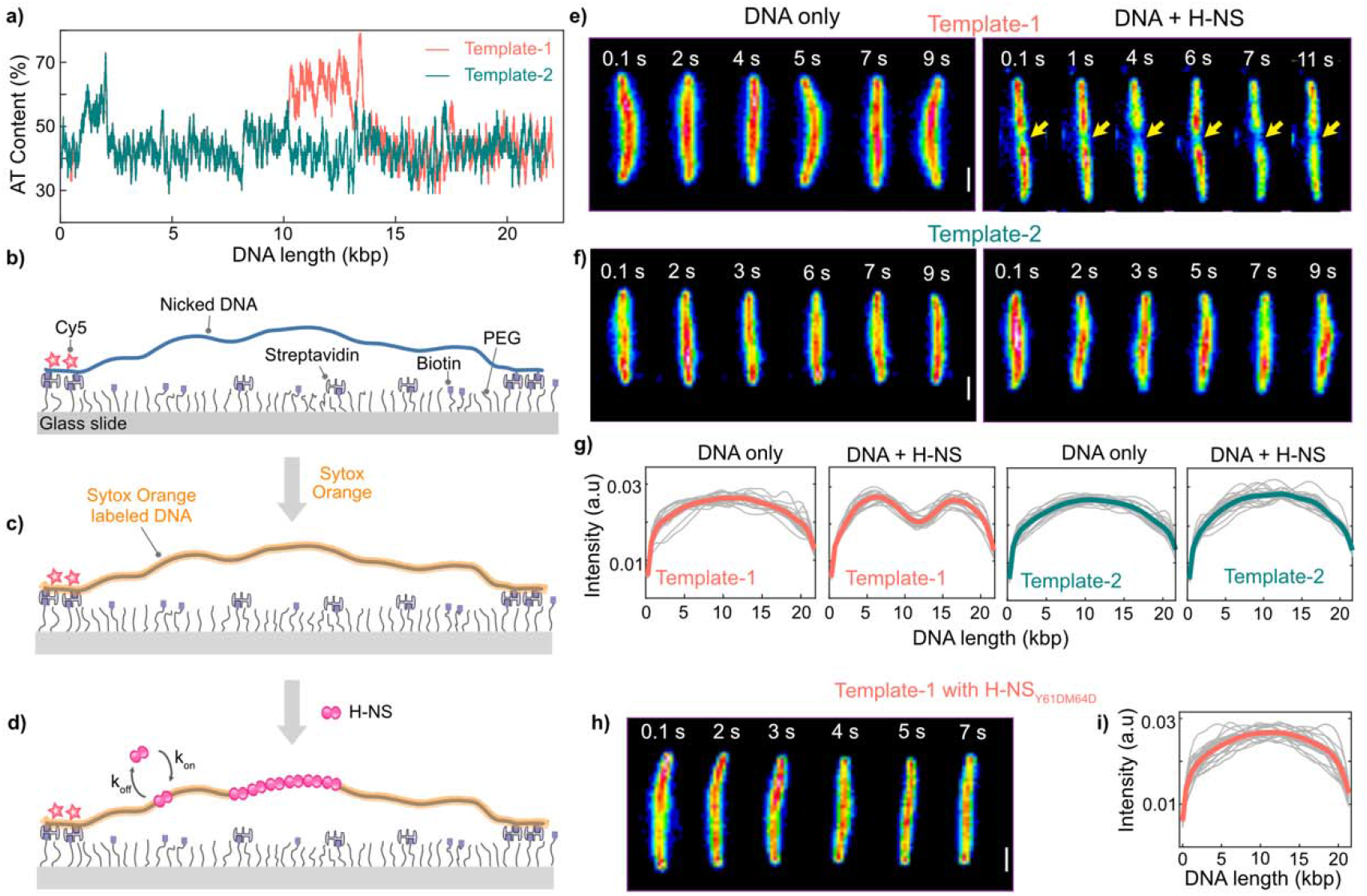
H-NS assembles sequence-specific nucleoprotein filaments at AT-rich loci on relaxed DNA. **(a)** Plots displaying the AT percentage along the DNA length for Template-1(Orange) with 3 kb 66% AT-rich site at the center and Template-2 (Dark green) with homogeneous AT-content. **(b)** Schematic of the single-molecule assay. A 21 kb DNA is immobilized at both ends via biotin-streptavidin interactions on glass surface. Cy5-label at one end marks the DNA direction. **(c)** The intercalating dye Sytox orange (SxO) is introduced into the channel which fluorescently stains DNA. **(d)** Schematic representation of H-NS (pink) bound DNA. **(e)** and **(f)** Representative snapshots of DNA without (left) and with H-NS (right) on corresponding Templates. The dark regions indicated with yellow arrow in the DNA+H-NS column for Tempate-1 shows filament formation at AT-rich loci. **(g)** Average fluorescence intensity profiles along normalized DNA contour (n ≥ 20 molecules per construct). **(h)** Representative snapshots of DNA with H-NS_Y61DM64D_. **(i)** Average fluorescence intensity profiles along normalized DNA contour (n ≥ 20 molecules per construct). Grey: individual molecules; Orange: mean for Template-1 and Green: mean for Template-2. Scale bar: 1μm (e, f, and h)

Upon introducing 500 nM wild-type H-NS, A distinct decrease in fluorescence intensity was observed precisely at the AT-rich region (Fig. 1e-right). This intensity depletion was stable throughout the imaging period of several minutes. Averaged fluorescence intensity profiles from many molecules showed a pronounced dip at the AT-rich locus in the presence of H-NS but a uniform profile without it (Fig. 1g). H-NS oligomerization on DNA manifests as a low intensity region along the homogeneously SxO-stained DNA contour, where the bound nucleoprotein filament excludes dye intercalation (Fig. 1e-right). We avoided fluorescent labeling of H-NS in all experiments, as such labels are known to perturb DNA-binding affinity and oligomeric state^13^; the intercalation-exclusion strategy provides a non-perturbative readout of H-NS filament formation.

We tested the sequence requirements for filament assembly on three additional constructs: Template-2 (homogeneous AT-content throughout), Template-3 (1 kb, 90% AT-rich central insert), Template-4 (a 100 bp high-affinity H-NS binding sequence^24^ in the center) and Template-5 (multi-AT-loci construct; Fig. 1a and Fig. S1a). While Template-2 showed no intensity heterogeneity upon H-NS addition (Fig. 1f and 1g), Template-3 and -4 produced pronounced dips in the fluorescence intensity in the center as observed in Template-1 (Fig. S1b and S1c), demonstrating that a contiguous high-AT-content or a high-affinity sequence seed is necessary for stable filament nucleation. Template-3 and Template-4, despite their difference in AT-tract length, produced fluorescence depleted regions of similar extent (Fig. S1d), indicating nucleation-limited, cooperative nucleoprotein filament propagation. Template-5 produced multiple fluorescence depletion regions at each of the AT-rich zones, confirming that filament nucleation occurs at each AT-rich zone independently of its position within the molecule (Fig. S1b and S1c).

The oligomerization-deficient double mutant H-NS_Y61DM64D_^13,25^, in which substitution of Tyr61 and Met64 with Asp amino acid disrupts the second dimerization interface, produced no dark regions on Template-1 and Template-5, in contrast to wild-type H-NS (Fig. 1i and S10). This confirms that the observed fluorescence exclusion is specifically reporting H-NS cooperative polymerization, not isolated dimer binding, and establishes a genetic tool for dissecting the two binding modes in subsequent experiments.

### H-NS filament acts as a topological barrier on positively supercoiled DNA

We next wanted to examine the effect of H-NS/DNA nucleoprotein filament on the dynamics of supercoiled DNA. For this, we generated positive DNA supercoiling *in situ* by introducing SxO to pre-tethered, topologically constrained molecules^21,22^ (Fig. 2a-2c). Intercalation locally unwinds the helix; because both tether ends are fixed, this introduces compensatory positive supercoiling globally, manifesting as plectonemic structures visible as bright fluorescent puncta.On bare positively supercoiled DNA, plectonemes diffused dynamically along the full contour of all DNA templates (Fig. 2d-2f and Fig. S2, Movie S2) with a mean diffusion coefficient D=12.88 ± 7.0 kbp^2^/s (Template-1) and 15.86 ± 7.6 kbp^2^/s (Template-2) (Fig. S2, Movie S6), consistent with previously published values for plectoneme diffusion^21^.

**Figure 2.**
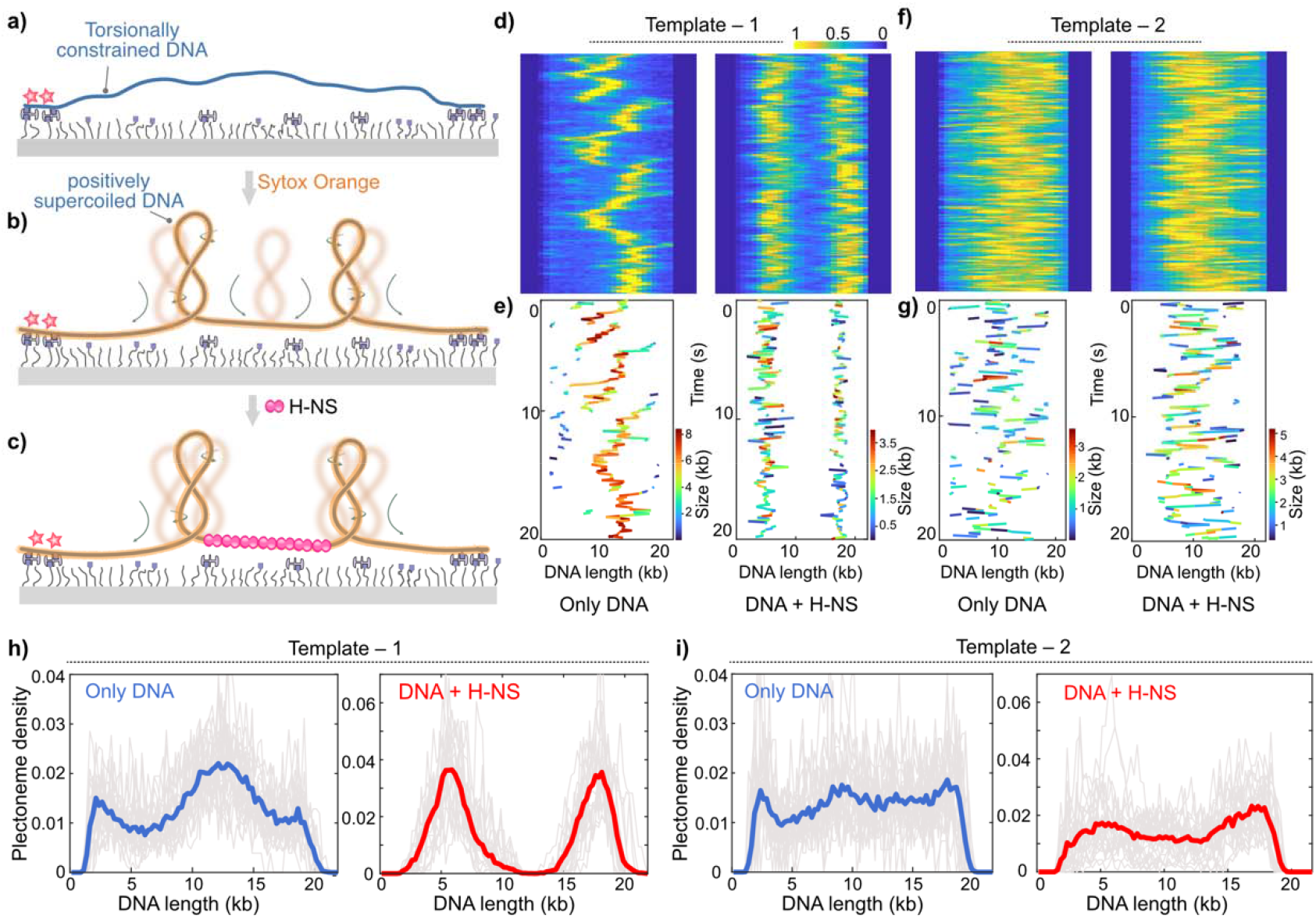
H-NS oligomerization on positively supercoiled DNA promotes topological barrier activity. **(a)** Schematic of DNA immobilization at both ends on a PEG-passivated surface and extended under laminar flow. **(b)** Intercalation of SxO generates positive supercoiling on a torsionally constrained DNA. Plectonemes diffuse freely along bare positively supercoiled DNA. **(c)** H-NS forms a filament and confines plectoneme diffusion to flanking regions. **(d)** Representative kymographs for Template-1 without (left) and with H-NS (right). **(e)** Plectoneme size along each frame corresponding to figure-(d). **(f)** Representative kymographs for Template-2 without (left) and with H-NS (right). **(g)** Plectoneme size along each frame corresponding to figure-(f). **(h)** Average plectoneme density profiles (1,000 frames; n ≥ 20 molecules) for Template-1 without (left) and with H-NS (right). **(i)** Average plectoneme density profiles (1,000 frames; n ≥ 20 molecules) for Template-2 without (left) and with H-NS (right). Scale bar: Horizontal scale – 2 μm (d and f) and vertical scale – 2 s (d and f).

Upon introducing H-NS onto positively supercoiled Template-1, a fluorescence depleted region appeared at the AT-rich location, indicating that H-NS adopts its filament mode even on supercoiled DNA. Strikingly, plectoneme diffusion became spatially restricted to the flanking regions and fully excluded from the H-NS filament zone (Fig. 2d-right 2e-right, Movie S3). Plectoneme density profiles derived from averages of several molecules each over 1,000-frame confirmed a Clear depletion of plectoneme occupancy at the H-NS filament locus (Fig. 2h and Fig. S2f). The mean diffusion coefficient on H-NS-bound DNA decreased to D = 3.72 ± 1.41 kbp^2^/s for Template-1(Fig. S2di), reflecting the reduction in accessible contour length. Notably, on bare positively supercoiled Template-1 and Tempate-3, plectoneme density at the AT-rich region was higher than in the flanking regions (Fig 2d-left, 2e-left, 2h-left and Fig S2e), indicating that this sequence is intrinsically favorable for plectoneme formation. H-NS binding completely reverses this preference, converting a plectoneme-enriched site into a plectoneme-depleted zone. On Template-2, no such barrier effect was observed in the presence of H-NS (Fig. 2f, 2g, and 2i, and Fig. S2, Movie S7), confirming that topological barrier formation requires sequence-specific filament nucleation.

The nucleoprotein acting as a barrier is physically intuitive; bending DNA at the rigid filament zone is energetically costlier than in the flanking flexible regions. As a result, plectonemes are effectively excluded from the filament and confined to the AT-poor regions on either side. The filament thus acts as a topological barrier, partitioning the supercoiled molecule into isolated domains on either flank. This demonstrates that H-NS filaments at AT-rich loci act as torsional boundaries, directly restricting supercoiling propagation and confining topological fluctuations to defined “chromosomal” regions.

### H-NS entirely switches into bridging mode on negatively supercoiled DNA

The physiologically predominant form of bacterial DNA is negatively supercoiled. To recapitulate this in our assay, we first incubated DNA molecules with 500 nM SxO prior to tethering. At this high concentration, SxO binds with high occupancy, each intercalation event locally unwinding the helix. Since the DNA ends are free at this stage, this unwinding is not compensated by supercoiling. After tethering both ends to the surface, the dye concentration was diluted to 100 nM. As intercalated dye molecules dissociate and re-equilibrate with the diluted solution, the torsional constraint imposed by the fixed ends causes the DNA to become underwound, generating negatively supercoiled plectonemes (Fig. 3a-3c). On bare negatively supercoiled DNA, plectonemes diffused freely along all templates with no obvious preferential localization (Fig. 3d-3g-left), similar to the behavior on positively supercoiled substrates.

**Figure 3.**
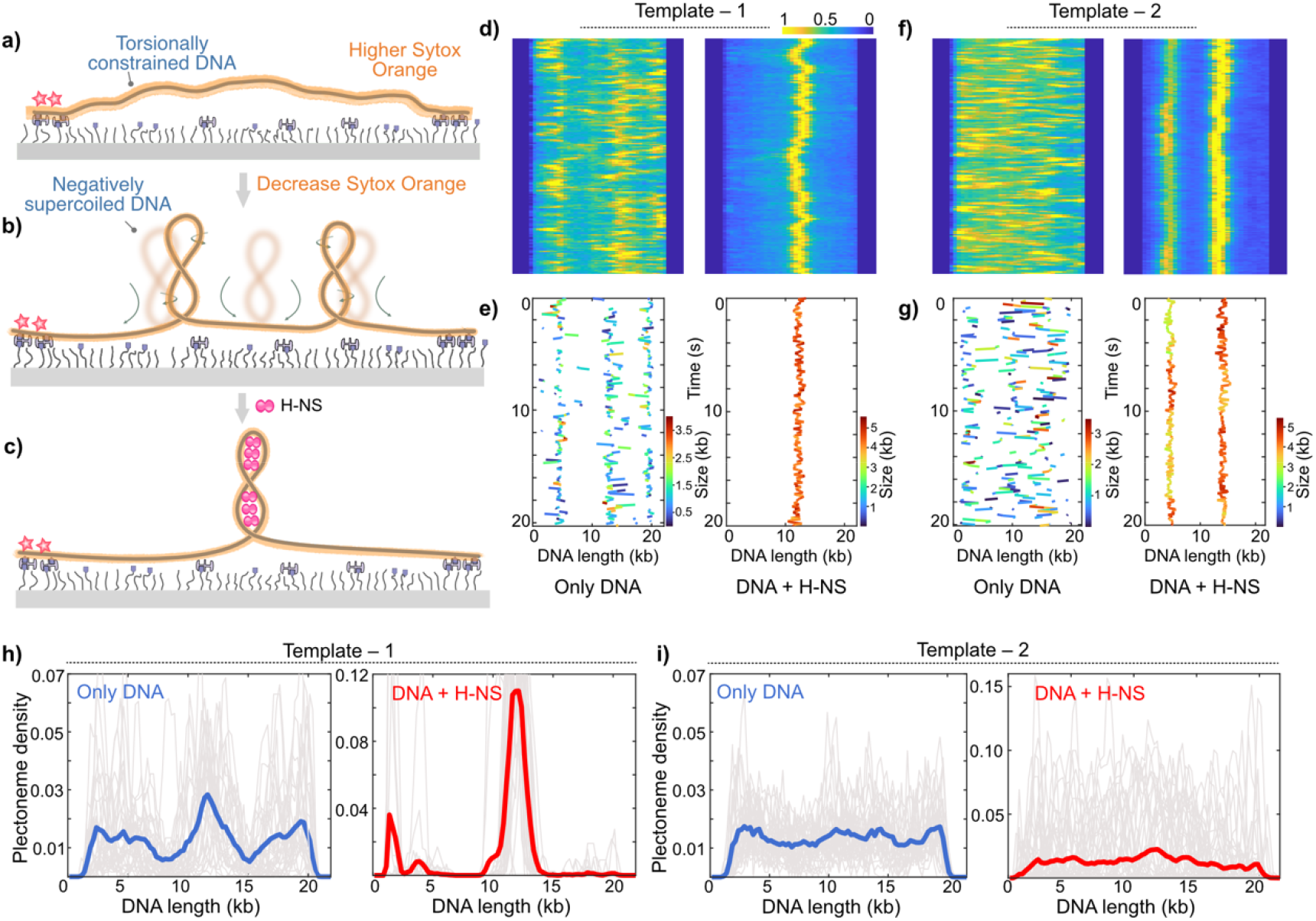
H-NS binds in bridging mode on negatively supercoiled DNA. **(a)** Schematic of DNA immobilization at both ends with high SxO concentration. **(b)** Decreasing SxO concentration generates negative supercoils. Plectonemes diffuse freely on bare negatively supercoiled DNA. **(c)** H-NS immobilizes a plectoneme as a stable high-intensity punctum at the AT-rich locus, indicating bridging mode. **(d)** Representative kymographs for Template-1 without (left) and with H-NS (right). **(e)** Plectoneme size along each frame corresponding to figure-(d). **(f)** Representative kymographs for Template-2 without (left) and with H-NS (right). **(g)** Plectoneme size along each frame for 10 seconds corresponding to figure (f). **(h)** Average plectoneme density profiles (1,000 frames; n ≥ 20 molecules) for Template-1 without (left) and with H-NS (right). **(i)** Average plectoneme density profiles (1,000 frames; n ≥ 20 molecules) for Template-2 without (left) and with H-NS (right). Scale bar: Horizontal scale – 2μm (d and f) and vertical scale – 2 s (d and f).

The response of H-NS to negative supercoiling was starkly different from its behavior on overwound or relaxed DNA. On negatively supercoiled Template-1 and Template-3, plectoneme density profile showed a weak preference for AT-rich region on bare DNA (Fig. 3h-left and Fig. S3e). However, H-NS produced *no dark region*, indicating absence of filament formation. Instead, we observed predominantly single static plectonemes as persistent, high-intensity puncta precisely at the AT-rich locus (Fig. 3d-3e-right, Fig. S3f, Movie S4, S9). Plectoneme density profiles showed a clear preference for AT-rich region (Figure 3h-right). This is the signature of bridging mode where H-NS cross-links the two juxtaposed DNA strands within the plectoneme, stabilizing it against diffusion. The mean diffusion coefficient of plectonemes dropped to D = 0.23 ± 0.3 kbp^2^/s, a ∼15-fold reduction relative to bare DNA (Fig. S3), demonstrating near-complete immobilization. Similar bridging on negatively supercoiled DNA was also observed upon including 1 mM Mg^2+^ in the buffer, in contrast to previous reports that low Mg^2+^ promote stiffening of H-NS^8^ (Fig. S7).

We further note that on negatively supercoiled Template-2 (homogeneous AT-content) and Template-4 (100 bp high affinity sequence of H-NS), H-NS could also stabilize plectonemes, but without any preferential localization showing uniform plectoneme density distribution (Fig. 3f, 3g, and 3i-right, Movie S8). Both the templates lack any preference for plectoneme formation on bare DNA (Fig. 3f, 3g, and 3i-left and Fig. S4). This reveals that while AT-richness biases the capture site by providing a preferred nucleation point, bridging itself exploits plectonemic geometry rather than sequence per se.

The mechanistic basis for this helicity-driven mode switch lies in the geometry of negatively supercoiled DNA. H-NS is a minor-groove binder, and the plectonemic superhelix brings two DNA duplexes into close apposition – within the ∼2–4 nm bridging span accessible to an H-NS dimer^11^ – providing a geometrically optimal substrate for cross-linking. The widened minor groove due to negative supercoiling and plectoneme-mediated proximity thus create conditions uniquely favorable for bridging on underwound DNA, conditions that do not exist on relaxed or overwound substrates where plectonemes are absent or where the minor groove geometry differs. AT-rich sequence and negative supercoiling are therefore cooperating inputs: sequence biases where bridging occurs, but helicity determines which mode operates. The fact that negative supercoiling selectively activates the bridging mode thus establishes DNA underwinding as the physiological trigger for H-NS-mediated gene silencing.

### H-NS senses changes in DNA helicity and switches binding modes in real-time

The opposing behaviors on positively and negatively supercoiled DNA raise the question of whether H-NS can respond dynamically to topological transitions on a physiologically relevant timescale. We addressed this with two complementary real-time switching experiments. First, starting from H-NS in bridging mode on negatively supercoiled Template-1 (Fig. 4a), upon nicking, the punctum intensity decreased progressively as the DNA relaxed towards a topologically unconstrained state (Fig. 4b). The plectoneme gradually disappeared in few seconds and was replaced by a fluorescence depleted region at the AT-rich locus (Fig. 4c and Fig. S9) – the hallmark signature of H-NS filament formation. The kinetics of the intensity transition is slower than the rate of supercoiled DNA relaxation,^26,27^ indicating that the topological change itself drives the mode switch and that H-NS reorganization is rate-limiting. This experiment directly demonstrates that the same H-NS molecules that were bridging on underwound DNA can reorganize into a stiffening filament within seconds as the DNA becomes relaxed.

**Figure 4.**
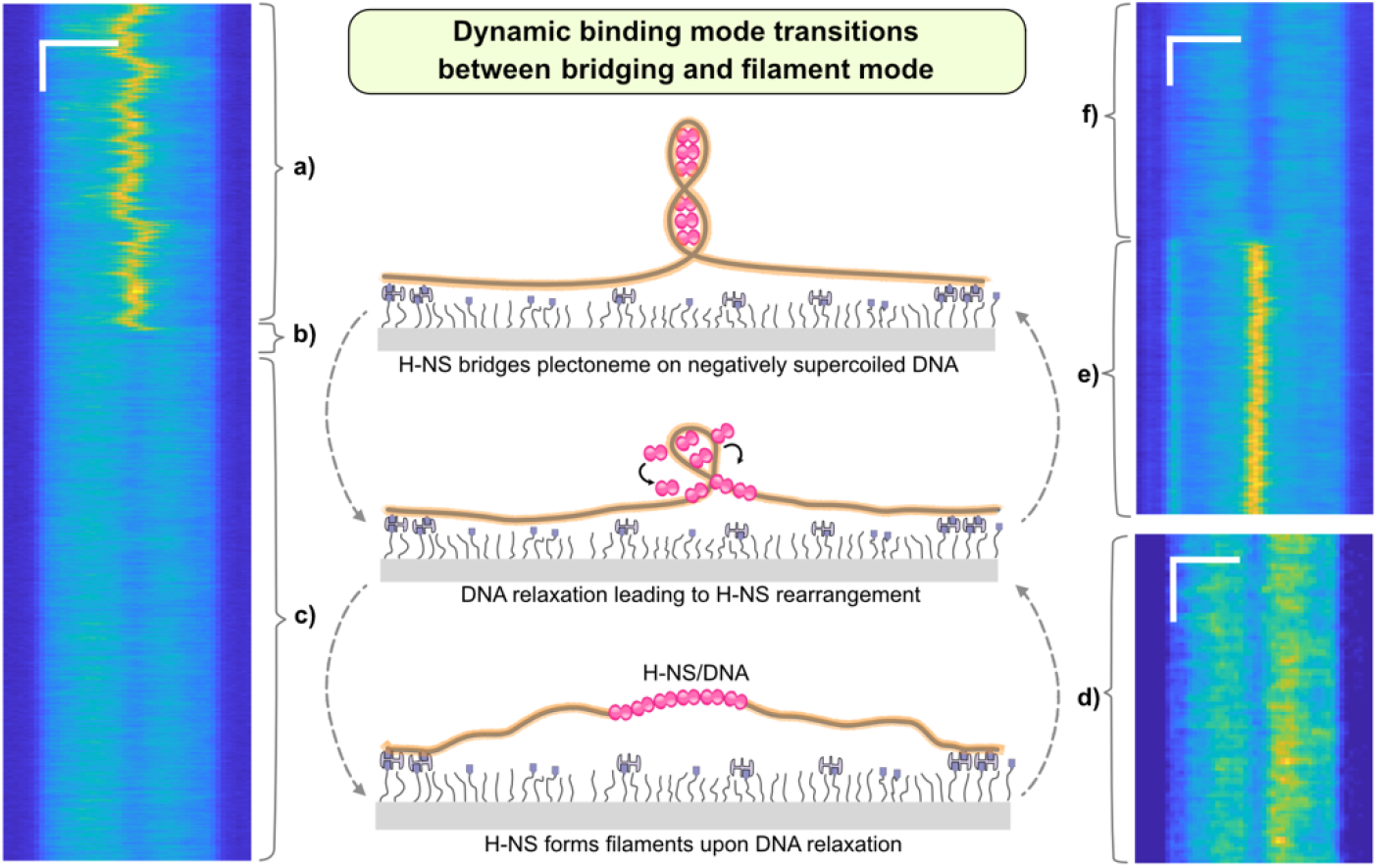
H-NS senses changes in DNA helicity and switches binding modes dynamically. **(a-c)** Bridging-to-stiffening switch triggered upon DNA relaxation from negative supercoiling for Template-1. Kymograph represents **(a)** H-NS in bridging mode: stable plectoneme punctum at AT-rich locus under negative supercoiling. **(b)** DNA nicking releases torsional constraint; punctum intensity decreases as DNA relaxes. **(c)** Relaxation complete; dark region replaces the punctum, reporting filament formation. **(d–e)** Stiffening-to-bridging switch triggered converting relaxed to negative supercoiling. Kymograph represents **(d)** Stiffening filament manifested by fluorescence depleted region in the center on relaxed Template-1 at high SxO. **(e)** SxO dilution induces negative supercoiling; fluorescence depleted region is replaced by a bridging punctum. **(f)** DNA nicking relaxes the DNA while maintaining H-NS nucleoprotein filament on the DNA. Scale bar: Horizontal scale – 2μm (a-f) and vertical scale – 2 s (a-f).

Second, we confirmed the reverse transition by an orthogonal approach. H-NS pre-incubated with relaxed DNA at high SxO concentration produced the dark-region signature of stiffening filaments upon immobilization. After diluting SxO to induce negative supercoiling, the dark regions were replaced by high-intensity bridging puncta (Fig. 4d–e), confirming stiffening-to-bridging conversion. Both transition directions were reproducible across all molecules in independent experiments. The switching timescale, on the order of seconds, is commensurate with the rate of supercoiling generation during transcription elongation^28^, making it feasible for the supercoiling wave from a single elongating RNA polymerase to toggle H-NS binding mode at flanking genes in real time. Similarly, on positively supercoiled DNA, we observed that H-NS filament acting as barrier for plectoneme diffusion became stable filament upon relaxation, and *vice versa* (Fig. S8a-S8d).

To confirm that mode switching reflects reorganization of stably bound H-NS rather than dissociation and rebinding of free protein, we performed washing experiments in which excess H-NS was removed from solution after filament formation on relaxed DNA. Even in the absence of free protein in solution, the bound H-NS responded to topological changes, inducing negative supercoiling converted the filament into a bridging punctum (Fig. S8g-S8j), while inducing positive supercoiling caused the bound protein to act as a barrier to plectoneme diffusion (Fig. S8e and S8f). This demonstrates that helicity-driven mode switching is an intrinsic property of chromosome-associated H-NS and does not require exchange with a solution pool of free protein, consistent with the behavior expected of a constitutively chromosome-bound architectural protein *in vivo*.

### The oligomerization-deficient H-NS_Y61DM64D_ mutant bridges but cannot switch to filament mode

To genetically dissect the two modes (filament and bridging) and confirm their mechanistic independence, we characterized the oligomerization-deficient double mutant H-NS_Y61DM64D_ on both supercoiled substrates. The Y61D/M64D substitutions disrupt the second dimerization interface in the H-NS N-terminal domain, impairing cooperative polymerization without affecting DNA binding^13,25^. On positively supercoiled Template-1, H-NS_Y61DM64D_ produced neither a dark region nor a plectoneme barrier (Fig. 5a), and density profiles showed no AT-locus depletion, only subtle position-independent modulation consistent with transient non-cooperative binding (Fig. 5d). Additionally, we observed a plectoneme mean diffusion coefficient (D ≈ 16.89 ± 9.36 kbp^2^/s; Fig.S6), similar to that on bare DNA. On negatively supercoiled Template-1, H-NS_Y61DM64D_ stabilized plectonemes at the AT-rich locus in a manner indistinguishable from wild-type where we observed persistent bridging puncta and plectoneme density profiles showed strong AT-locus enrichment (Fig. 5b, 5c, and 5e), and the mean diffusion coefficient was near-zero (D ≈ 0.58 ± 0.16 kbp^2^/s; S6). This confirms that H-NS bridging requires only dimeric protein and that the second dimerization interface is dispensable for bridging, consistent with a two-pin bridging architecture in which a single H-NS dimer cross-links two duplexes^11^. Upon nicking of H-NS_Y61DM64D_-bound negatively supercoiled DNA, the bridging punctum dissipated as the DNA relaxed, but no fluorescence depleted region appeared (Fig. 5c). The mutant sensed the topological change and released from bridging plectoneme but could not reorganize into a filament. Together, these results confirm that the two modes are mechanistically distinct and can be uncoupled: an H-NS dimer is sufficient for bridging but stiffening requires oligomerization of dimers^13,25^, and our data demonstrate that DNA helicity is the switch that selects between them.

**Figure 5.**
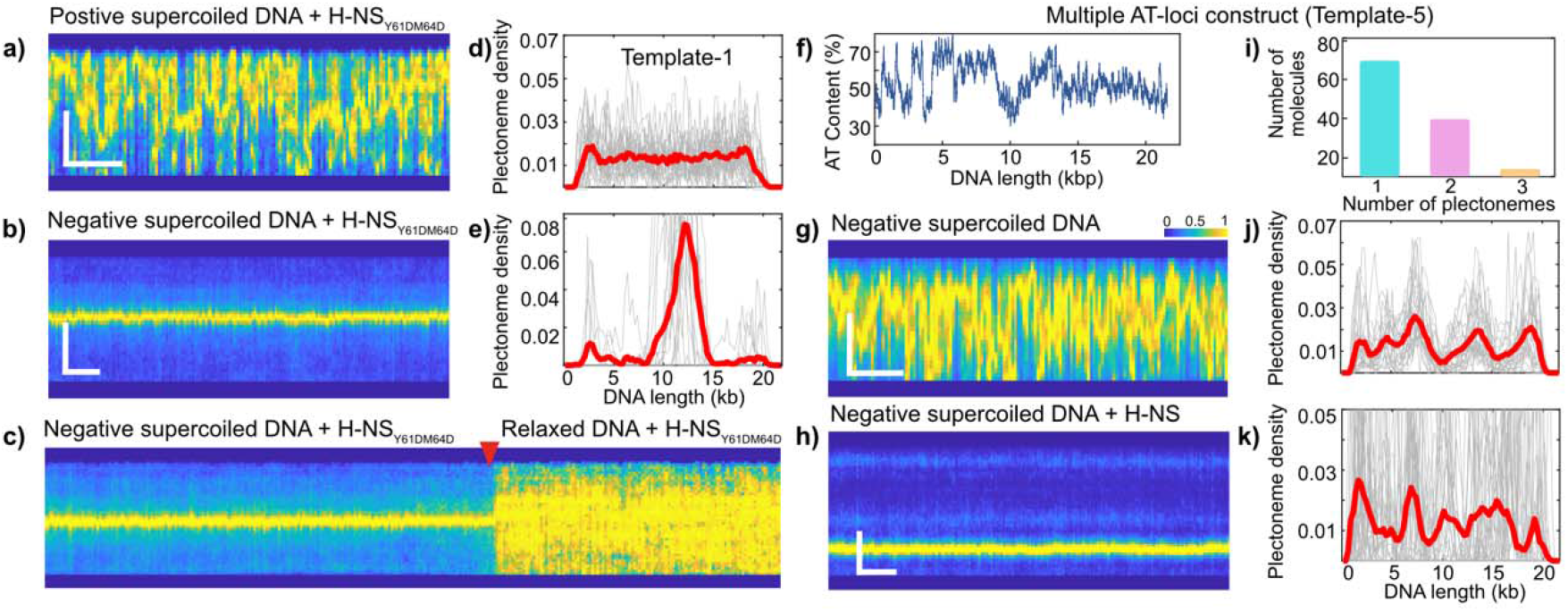
The oligomerization-deficient H-NS_Y61DM64D_ mutant bridges while helicity-driven mode selection yields self-organized silencing and insulation domains on multi-locus DNA. **(a)** and **(b)** Representative kymographs of positively and negatively supercoiled DNA, respectively, in the presence of H-NS_Y61DM64D_. **(c)** Kymograph showing transition (Red arrow) from bridged punctum on negatively supercoiled DNA to homogenous intensity upon DNA relaxation. **(d)** and **(e)** Average plectoneme density profile for positive supercoiled DNA and negative supercoiled DNA, respectively, with mutant H-NS. **(f-o)** Behavior of H-NS on multi-AT locus DNA. **(f)** Plot displaying AT-content for Template-5 along the DNA length. **(g)** and **(h)** Representative fluorescence kymographs of Template-5 under negative supercoiled DNA without H-NS and with H-NS, respectively. **(i)** Plot displaying the number of plectonemes bridged on each molecule on Template-5. **(j)** and **(k)** Average plectoneme density profiles of Template-5 over the 1000 frames for ≥ 20 molecules for negative supercoiled DNA without H-NS and with H-NS, respectively. Scale bar: Horizontal scale – 2μm (a-h) and vertical scale – 2 s (a-h).

### Helicity-driven mode selection drives self-organized chromosomal domain formation

Having established that DNA helicity governs H-NS binding mode at individual AT-rich loci, we asked what consequence this mode-switching mechanism produces at the chromosomal scale, where multiple AT-rich loci are distributed along a single topologically constrained DNA. To address this, we utilized a 21 kb DNA bearing largely AT-rich segments, separated by GC-rich spacers (Fig. 5f). Under negative supercoiling, on bare DNA, plectonemes once again showed dynamic diffusion on Template-5 (Fig. 5g and 5j). We then introduced H-NS and imaged the resulting organization.

Strikingly, a subset of AT-rich sites displayed a high-intensity bridging punctum while the remaining showed fluorescence depleted regions characteristic of nucleoprotein filaments (Fig. 5h-5k, Fig. S9a). Bridging and filament formation coexisted simultaneously on the same molecule and at the same time, directly demonstrating the complementarity between the two H-NS binding modes. Those sites that capture the available plectonemes enter bridging mode compatible for their transcriptional silencing, while flanking sites adopt filament mode to insulate the resulting domain from neighboring torsional fluctuations. Both topological outputs required for silencing and insulation emerge from the same helicity-sensing mechanism operating at different loci under different local plectoneme occupancy.

Plectoneme density plot for negatively supercoiled DNA showed specific, reproducible pattern across molecules (Fig. 5j). H-NS binding led to a pattern that is correlated with bare DNA density (Fig. 5k), but the bridging sites varied stochastically across molecules (Fig. S9a), and each segment could serve as the bridging site. This stochasticity reflects that all AT-rich sites are competent for plectoneme capture, and whichever site the diffusing plectoneme first encounters locks in as the bridging site while the others become filament sites. Interestingly, under H-NS bridging, the plectonemes are not static but exhibit slow dynamic rearrangement, characterized by the shrinkage of an existing bridged plectoneme and the simultaneous growth of another at a distant site (Fig. S9b-S9f). In a clonal bacterial population, this mechanism predicts that different cells will silence different genes despite identical DNA sequence and identical H-NS levels, a physical topology-driven source of cell-to-cell heterogeneity in gene silencing that requires no dedicated regulatory circuitry^29,30^. The oligomerization mutant H-NS_Y61DM64D_ on the multi-AT-loci construct produced plectoneme pinning without flanking fluorescence depleted regions (Fig. S11), confirming that the insulating filaments at non-bridging sites require oligomerization and are genetically separable from the bridging event itself.

## Discussion

The central finding of our work is that DNA superhelical polarity is the physiological determinant of H-NS binding mode selection. On positively supercoiled DNA, H-NS polymerizes into stiffening filaments; on negatively supercoiled DNA, H-NS bridges apposed duplexes. This establishes a direct physical link between the dynamic torsional state of the bacterial chromosome and the functional output of its most abundant gene-silencing protein. The mechanistic logic of this helicity-sensing switch is grounded in the structural properties of the two substrates, positively and negatively supercoiled DNA. On positively supercoiled DNA, the plectoneme geometry may not favor H-NS bridging. One possibility is that the two strands do not adopt the inter-duplex spacing or minor groove width required for cross-linking, though we cannot directly resolve this from our data. Instead, H-NS appears to polymerize along the single duplex in stiffening mode, exploiting the AT-rich minor groove as a template for cooperative protein assembly. On negatively supercoiled DNA, the widening of the minor groove and plectoneme-mediated duplex proximity may collectively create a binding geometry favorable for bridging, consistent with our single-molecule observations, though the precise structural basis of this preference remains to be established. The two modes are therefore responses to structurally distinct substrates that happen to be generated by opposite polarities of torsional stress in the same molecule.

This interpretation resolves the long-standing debate over ionic control of H-NS mode switching. Prior *in vitro* work identified elevated magnesium concentration as a promoter of bridging, proposing it as a physiological trigger^12,13^. However, intracellular free Mg^2+^ (∼1–2 mM)^31-33^ falls well below the concentrations required (above 5 mM) for magnesium-induced bridging in those assays, an inconsistency that has undermined confidence in ionic control as a physiological mechanism. Our experiments at physiological magnesium concentrations demonstrate robust, complete, and reversible mode switching driven purely by superhelical polarity. We suggest that topology is the dominant *in vivo* switch rather than elevated divalent cation concentration because DNA supercoiling is dynamic and directly coupled to transcriptional activity.

The connection between our findings and transcription is particularly significant considering the twin-supercoiling domain model, according to which an elongating RNA polymerase generates negative supercoiling upstream and positive supercoiling downstream of the transcription bubble^19^. Our data predict that H-NS immediately downstream of an elongating polymerase would adopt filament mode because of positive supercoiling, creating a torsional barrier that prevents it from propagating into flanking regions. Upstream, H-NS would bridge negative supercoil plectonemes at AT-rich loci, acting as a molecular clutch and compete with the transcription machinery for the same torsional resource. Rather than functioning as a simple on/off silencer, H-NS may thus fine-tune transcriptional output by modulating the availability of transcription-facilitating negative supercoiling.

The multi-AT-locus experiment reveals the chromosomal-scale consequence of helicity-driven mode selection. The two H-NS binding modes serve spatially distinct functions — bridging to silence at one site while filaments insulate at flanking sites. This self-organized pattern means that H-NS does not need dedicated domain boundary sequences or additional regulatory factors to organize the chromosome. The topology-sensing property inherent to the protein itself, combined with the physical constraints of a topologically closed DNA, is sufficient to generate a complete, insulated gene-silencing domain. The stochasticity of bridging site selection in the multi-locus system has consequences for bacterial cell biology that extend beyond chromosome organization. H-NS-silenced genes—including virulence factors in *Salmonella*, metabolic islands in *E. coli*, and sporulation regulators in *Bacillus subtilis*—are known to exhibit heterogeneous, noisy expression across isogenic populations, contributing to population-level phenotypic heterogeneity that buffers against fluctuating or stressful conditions^29,30^. Stochastic competition among AT-rich loci for the available negative supercoiling generates different silencing configurations across cells without requiring any mutation, epigenetic mark, or dedicated regulatory circuit. The same mechanism that organizes the chromosome therefore also generates adaptive phenotypic diversity across the population.

Our findings provide a mechanistic framework for interpreting recent genome-wide observations of H-NS-dependent chromosome organization. Ultra-high-resolution Micro-C in *E. coli* revealed that H-NS and StpA organize the nucleoid into discrete chromosomal hairpins and hairpin domains that directly repress horizontally transferred genes, with disruption of H-NS causing extensive local reorganization of the 3D genome^4^. Our single-molecule data suggest that these structural elements are the population-level readout of the helicity-driven bridging events we describe here, occurring at AT-rich loci under the predominant global negative supercoiling *in vivo*. Separately, Hi-C in stationary-phase *E. coli* demonstrated that H-NS mediates large-scale DNA bridging that reshapes the transcriptional landscape as cells exit exponential growth^5^, consistent with our observation that bridging mode is the dominant mode under physiological negative supercoiling. Notably, transcription has been shown to counteract H-NS silencing at a distance through supercoiling propagation rather than by direct polymerase encroachment on the silenced locus^17^ which our mechanism explains directly. Positive supercoiling generated by an upstream elongating polymerase propagates to the H-NS-bound locus and switches it from bridging to filament mode, releasing silencing without the polymerase ever reaching the gene.

DNA bridging is not unique to H-NS among NAPs — IHF, HU, Fis, and Dps all bridge or compact DNA, but do so in a topology-independent manner or through geometrically distinct mechanisms unrelated to plectoneme stabilization^3^. Similarly, Dps bridges DNA independently of both sequence and topology^34^. What distinguishes H-NS is therefore not bridging per se, but that its bridging is selectively activated by negative supercoiling, geometrically coupled to plectoneme formation, and reversibly switched off by positive supercoiling — making it uniquely suited to read and respond to the dynamic torsional landscape generated by transcription. Intriguingly, Lsr2, the functional xenogeneic silencer in *Mycobacterium*, compacts DNA through sequence-dependent co-condensation at AT-rich regions, but not forming filaments^35^. This contrast suggests that functional equivalence in xenogeneic silencing does not necessarily imply mechanistic equivalence in topology sensing. Notably, GapR from *Caulobacter crescentus* shows the opposite topology preference, associating preferentially with positively supercoiled DNA^36^, suggesting that supercoiling polarity sensing may be a broader organizing principle among DNA-binding proteins, with the preferred superhelical state tuned to the specific biological function of each protein. The single-molecule approach we describe here provides a direct and generalizable framework for investigating how NAPs read and respond to the dynamic torsional landscape of the bacterial chromosome.

## Acknowledgments

The authors used Claude (Anthropic) for assistance with manuscript editing. We would like to thank all the Ganji lab members for their meaningful discussions. This work has been supported in part by a DBT/Wellcome India Alliance intermediate fellowship (IA/I/21/2/505928) and the Department of Biotechnology (BT/PR40186/BTIS/137/3/2020). We greatly acknowledge the support from Max-Planck Institute of Immunology and Epigenetics Freiburg, Germany in terms of partner group to Mahipal Ganji lab.

## Author contributions

**SS:** Conceptualization, Methodology, Investigation, Visualization, Writing – original draft, review & editing; **SB:** Conceptualization, Methodology, Investigation, Visualization, Writing – review & editing; **SG:** Visualization; **MG:** Conceptualization, Methodology, Funding acquisition, Project administration, Supervision, Writing – original draft, review & editing

## Data and materials availability

All data are available in the main text or the supplementary materials. Raw data will be provided upon request.

