## Supplementary methods and figures for "A DNA-binding protein senses DNA superhelicity to switch between bridging and nucleoprotein filament formation"

### MATERIALS AND METHODS

#### DNA construct preparation.

#### Template-1: 21 kb DNA with 3 kb 66% AT-content insert at the centre.

A 21 kb DNA molecule harboring a 3 kb 66% AT-rich region at its center was prepared by insertion of the AT-rich fragment into an 18 kb vector backbone[1]. The 3 kb AT-rich region was PCR-amplified from Lambda phage DNA using gene-specific primers incorporating an XbaI site via the forward primer (3 kb 66% AT + XbaI_FP) and a PciI site via the reverse primer (3 kb 66% AT + PciI_RP) (Supplementary Table 1). The amplified fragment was digested with XbaI (NEB, R0145S) and PciI (NEB, R0655S) and ligated into the 18 kb plasmid backbone pre-digested with the same enzymes using T4 DNA ligase (NEB, M0202S). The resulting ligated product was transformed into chemically competent *E. coli* DH5α cells, and positive colonies harboring the 3 kb AT-rich insert were identified by colony PCR.

To generate biotinylated DNA molecules for single-molecule experiments, the 21 kb plasmid was digested with NotI and XhoI, followed by clean-up to remove non-specific small fragments. Biotinylated handles were prepared from PCR amplification of 500 bp fragments from pBluescript plasmid. The PCR reaction contains modified nucleotide (Biotin-11-dUTP, Jena Bioscience, NU-803-BIOX-S) along with the unmodified nucleotides. PCR-amplified DNA fragments containing multiple biotin moieties were ligated to each end of the linearized molecule using T4 DNA ligase. To confer directionality, Aminoallyl-dUTP-Cy5 (Jena Bioscience, NU-803-CY5-L) was incorporated into one of the 500 bp biotin handles. Specifically, the XhoI-digested end of the 21 kb molecule was ligated to an XhoI-digested 500 bp biotin handle, while the NotI-digested end was ligated to a NotI-digested 500 bp biotin handle.

The ligated product was resolved from non-ligated fragments by electrophoresis on a 0.8% agarose gel, stained with ethidium bromide, and visualized using a fluorescence gel documentation system (GelDoc). The 21 kb band was excised, and the DNA was extracted from the gel slice for subsequent use in single-molecule experiments. The biotinylated DNA molecules with end-specific Cy5 labeling were subsequently prepared following the same strategy as described in Template 1.

**Template 2: 21 kb DNA with 3 kb moderate AT-content as a control.** A 21 kb control DNA construct was prepared as follows. The 3 kb region of moderate AT-content (44%), comparable to that of the 18 kb backbone vector, was PCR-amplified from Supercos Lamda plasmid (pSuperCos-λ1,2 ) using forward primer (3 kb AT + XbaI_FP) incorporating an XbaI site and reverse primer (3 kb AT + PciI_RP) incorporating a PciI site (Supplementary Table 1). The amplified fragment was gel-extracted and digested with

PciI and XbaI restriction enzymes, and the digested product was ligated into the pre-digested 18 kb backbone vector. The resulting ligated product was transformed into chemically competent *E. coli* DH5α cells, and positive colonies harboring the 3 kb insert were identified by colony PCR and sequencing. Biotinylated DNA molecules with end-specific Cy5-labeling, including handle ligation, gel electrophoresis, band excision, and gel extraction, were subsequently prepared following the same strategy as described for Template 1.

**Template 3: 21 kb DNA with 1 kb 90% AT - rich insert at the center.** The Template-3 was prepared from 18 kb plasmid by incorporating 1 kb 90% AT rich site and was kindly provided by the laboratory of Prof. Krishanpal Karmodiya (IISER Pune). The plasmid was linearized by double digestion with NotI and XhoI restriction enzymes. Biotinylated DNA molecules with end-specific Cy5-labeling, including handle ligation, gel electrophoresis, band excision, and gel extraction, were subsequently prepared following the same strategy as described for Template 1.

**Template 4: 18 kb DNA with 100 bp H-NS nucleation site.** An 18 kb DNA molecule harboring a 100 bp H-NS nucleation site at its center was prepared by insertion of the 100 bp into an 18 kb vector backbone[1]. A 100 bp H-NS nucleation site fragment was a kind gift from Naganathan Lab, IIT-Madras. The 100 bp fragment was PCR amplified with using forward primer (H-NS_100bp_XbaI_FP) incorporating an XbaI site and reverse primer (H-NS_100bp_HindIII_RP) incorporating a HindIII site (Supplementary Table) The amplified fragment was digested with XbaI and HindIII (NEB, R3104S) and ligated into the 18 kb plasmid backbone pre-digested with the same enzymes. The resulting ligated product was transformed into chemically competent *E. coli* DH5α cells, and positive colonies harboring the 100 bp insert were identified by colony PCR and sequence verified. Biotinylated DNA molecules were subsequently prepared as described for Template 1.

**Template 5: Multi-AT-locus DNA.** Template 5 was prepared from the engineered plasmid pSuperCos-λ1,2, which was kindly provided by the laboratory of Prof. Cees Dekker (Delft University of Technology)[2]. The plasmid was linearized by double digestion with NotI and XhoI restriction enzymes. Biotinylated DNA molecules with end-specific Cy5-labeling, including handle ligation, gel electrophoresis, band excision, and gel extraction, were subsequently prepared following the same strategy as described for Template 1.

**Microfluidic channel preparation and DNA tethering**

Microfluidic flow cells were assembled from glass coverslips passivated by covalent PEG functionalization as described earlier (mPEG-SVA-5000/Biotin-PEG-SVA-5000),40:1 molar ratio)[3]. Prior to DNA immobilization, the microfluidic channels were subjected to a sequential surface blocking and functionalization protocol. Channels were first incubated with 40 µl of 0.1 mg/ml BSA (Sigma-Aldrich) for 5 min, followed by washing with

200 µl of T50 buffer (50 mM Tris-HCl pH 7.5, 50 mM NaCl). Channels were then incubated with 40 µl of 1% Tween 20 (Sigma-Aldrich) for 5 min and washed with 200 µl of T50 buffer. This was followed by incubation with 40 µl of 0.2 mg/ml neutravidin (Thermo Fisher) for 5 min and subsequent washing with 200 µl of T50 buffer to remove any unbound neutravidin.

For DNA tethering, 30 pM biotinylated DNA in T50 buffer was flushed through the channel at a flow rate of 15 µl/min, allowing hydrodynamic flow-induced extension of DNA molecules and sequential end-tethering to the neutravidin-functionalized surface. The channel was subsequently washed with 200 µl of T50 buffer to remove untethered DNA molecules prior to imaging.

**Single-molecule fluorescence imaging**

Single-molecule fluorescence imaging was performed on a Nikon Ti2 Eclipse microscope equipped with a motorized H-TIRF module and a perfect focus system to maintain focal stability throughout image acquisition. Illumination was provided by 561 nm and 640 nm wavelength lasers using the L6cc laser combiner (Oxxius Inc., France), and all imaging was performed under highly inclined and laminated optical sheet (HiLo) illumination conditions. An oil-immersion objective lens (Nikon Instruments Apo SR TIRF 100×, numerical aperture 1.49, oil) was used for all experiments. Fluorescence emission was detected using a Teledyne Photometrics PRIME BSI sCMOS camera operated at 16-bit readout sensitivity, with 2×2 pixels binning and cropped to an effective field of view of 1024×1024 pixels, corresponding to a pixel size of 130×130 nm and an effective imaging area of 130×130 µm at the sample plane.

Images were acquired at 100 msec exposure time per frame. Sequential dual-color imaging was performed by first illuminating with the 640 nm laser to localize Cy5 signals on DNA, followed by continuous 561 nm laser illumination to image SxO-stained DNA molecules. Kymographs were generated by extracting an 11-pixel-wide line profile along the DNA contour in each frame using published MATLAB scripts[2,4].

**Generation of positively and negatively supercoiled DNA**

Supercoiling of tethered DNA molecules was achieved by exploiting the helix-unwinding properties of the intercalating dye SYTOX Orange (SxO; S11368, Thermo Fisher), which induces compensatory supercoiling in topologically constrained DNA molecules[2,4]. All experiments were performed in an imaging buffer consisting of 50 mM Tris-HCl pH 7.5, 50 mM NaCl, 300 nM protocatechuate dioxygenase (PCD), 2 mM Trolox, and 2.5 mM protocatechuic acid (PCA) as an enzymatic oxygen scavenging system unless mentioned otherwise.

For positive supercoiling, biotinylated DNA molecules (30 pM) were immobilized in the microfluidic flow cell at a flow rate of 15 µl/min. Untethered DNA molecules were subsequently removed by washing with T50 buffer. Positive supercoiling was then induced by flowing 100 nM SxO in imaging buffer at 50 µl/min over the pre-tethered, topologically constrained DNA molecules. Intercalation of SxO locally unwinds the DNA helix, generating compensatory positive supercoiling in doubly tethered molecules, which was visualized by the appearance of plectoneme structures. Fluorescence images were acquired at 100 msec exposure time per frame, with initial illumination using the 640 nm laser for the first 100 frames for Cy5 localization, followed by continuous 561 nm laser illumination for imaging of SxO-stained DNA.

For negative supercoiling, biotinylated DNA molecules (30 pM) were first incubated with 500 nM SxO in imaging buffer in a tube for 2 min to achieve complete intercalator saturation. The SxO-saturated DNA was then immobilized in the microfluidic flow cell at a flow rate of 15 µl/min. Following immobilization, negative supercoiling was introduced by rapid buffer exchange with imaging buffer containing 100nM SxO, thereby reducing the intercalator concentration and causing re-winding of the DNA helix beyond its relaxed state in the topologically constrained molecules. Imaging conditions were identical to those described for positive supercoiling. Topological state was verified in both cases by monitoring plectoneme dynamics and fluorescence intensity profiles, which are characteristic indicators of the superhelices density and sign of supercoiling[2,4].

**Cloning, expression, and purification of H-NS**

Genomic DNA was extracted from *E. coli* BL21 (DE3) using the phenol-chloroform extraction method. The H-NS gene sequence was retrieved from the *E. coli* BL21 (DE3) complete genome database (NCBI), and gene-specific primers incorporating vector-compatible overlapping sequences for HiFi Assembly were designed such that the reverse primer encoded a C-terminal $His_6$ tag immediately upstream of the stop codon. The primers used were Ec H-NS FP (forward) and Ec H-NS RP (reverse), where the reverse primer was designed to incorporate a C-terminal 6×His-tag sequence followed by the stop codon.

The H-NS gene was PCR-amplified from *E. coli* BL21 (DE3) genomic DNA using the above primers. The pET28a (+) vector was linearized by PCR using primers flanking the insertion site. The H-NS PCR amplicon and the linearized pET28a (+) vector were assembled at a 1:2 molar ratio of vector to insert and assembled using the NEB HiFi DNA Assembly Master Mix (NEB, E2621). The assembly reaction was incubated at 50 °C for 60 min, after which the assembled product was transformed into chemically competent *E. coli* TOP10 cells. Transformants were selected on LB agar plates supplemented with 50 µg/ml kanamycin. Positive clones were initially screened by double restriction digestion with BamHI and PciI (New England Biolabs) to verify the presence and correct size of the H-

NS insert and subsequently confirmed by Sanger sequencing (Barcode Biosciences) to verify the correct insertion and reading frame of the H-NS gene. The sequence-verified construct, designated pET28a (+)-HNS-cHis6, was stored at −20 °C for subsequent use in protein expression.

H-NS protein was expressed in *E. coli* BL21 (DE3) cells harboring the pET28a (+)-HNS-cHis6 plasmid. Cultures were grown at 37 °C in Luria broth containing 50 µg/ml kanamycin until an $OD_{600}$ of 0.4 was reached. Protein expression was induced by reducing the temperature to 30°C and adding 500 µM IPTG, followed by continued growth for 4 hours while shaking at 180 rpm. Cells were harvested by centrifugation at 6,000×rpm for 45 min at 4 °C and resuspended in H-NS lysis buffer (20 mM Tris–HCl pH 7.5, 100 mM NaCl, 5% glycerol, 2 mM EDTA, and 1 mM DTT) containing 0.1 mg/ml PMSF. Cells were lysed by sonication for 30 min at 37% amplitude with 5 sec ON and 10 sec OFF pulse cycles. The lysate was clarified by centrifugation at 11,000×g for 45 min at 4 °C, and the resulting supernatant was subjected to polyethyleneimine (PEI; average molecular weight of 60 kDa; Sigma-Aldrich) precipitation by adding PEI to a final concentration of 0.6% with 15 min incubation, and the H-NS-containing precipitate was recovered by centrifugation at 11,000×g for 45 min. The pellet was washed by gentle resuspension in PEI wash buffer (10 mM Tris–HCl pH 7.5, 150 mM NaCl, 0.1 mM EDTA, 5% glycerol, and 1 mM DTT) and centrifuged at 11,000×g for 45 min. H-NS was subsequently eluted by gentle resuspension in PEI elution buffer (10 mM Tris–HCl pH 7.5, 600 mM NaCl, 0.1 mM EDTA, 5% glycerol, and 1 mM DTT), followed by centrifugation at 11,000×g for 45 min. The resulting supernatant was subjected to ammonium sulfate precipitation by slow addition with gentle stirring to a final concentration of 70%, and stirring was continued until the ammonium sulfate was completely dissolved, and the precipitate was recovered by centrifugation at 27,000×g for 45 min.

The H-NS containing precipitate was resuspended in buffer A (20 mM Tris–HCl pH 7.5, 500 mM NaCl, and 1 mM DTT) supplemented with 5 mM imidazole and applied to a Ni-NTA affinity column. The column was washed with buffer A containing 5 mM imidazole, and bound protein was eluted using a linear gradient of 10–500 mM imidazole. Fractions containing H-NS were pooled and dialyzed overnight against buffer B (10 mM Tris–HCl pH 7.5, 0.1 mM EDTA, 5% glycerol, 100 mM NaCl, and 1 mM DTT). The dialyzed protein was applied to a heparin column pre-equilibrated with buffer B, washed extensively, and eluted with a linear gradient of 0.1–0.9 M NaCl. H-NS containing fractions were pooled and dialyzed into storage buffer (20 mM Tris–HCl pH 7.5, 300 mM NaCl, and 10% glycerol), snap-frozen in liquid nitrogen, and stored at −80 °C until further use.

**Cloning, expression, and purification of H-NS$_{Y61DM64D}$**

The H-NS Y61D/M64D double mutant was generated by sequential introduction of mutations into the wild-type pET28a (+)-HNS-cHis$_6$ construct using a primer-based mutagenesis approach combined with NEB HiFi DNA assembly. The Y61D mutation was first introduced by PCR amplification of the H-NS gene using mutagenic primers encoding the Y61D substitution, designated Ec H-NS Y61D FP (forward) and Ec H-NS Y61D RP (reverse) (Table S1). The assembly, transformation, and clone verification were performed as described for wild-type H-NS cloning. The sequence-verified construct, designated pET28a (+)-H-NS-Y61D-cHis$_6$, was subsequently used as the template for introduction of the second mutation. The M64D mutation was then introduced into the pET28a (+)-HNS-Y61D-cHis$_6$ construct following the same primer-based HiFi assembly approach, using mutagenic primers designated Ec H-NS M64D FP (forward) and Ec H-NS M64D RP (reverse) (Table S1).

The assembly, transformation, and clone verification were performed as described for wild-type H-NS cloning. The sequence-verified double mutant construct, designated pET28a (+)-HNS-Y61DM64D-cHis6, was stored at −20 °C for subsequent use in protein expression.

Expression and purification of H-NS Y61DM64D was performed following a modified protocol from wild-type H-NS, including overexpression in *E. coli* BL21 (DE3), Ni-NTA affinity chromatography, and heparin column chromatography. The PEI precipitation and ammonium sulfate precipitation steps were omitted for the mutant purification, as no visible precipitation pellet was observed for H-NS Y61DM64D during the PEI step. This is likely due to the impaired ability of the Y61DM64D mutant to oligomerize onto DNA, since PEI precipitation relies on the co-precipitation of nucleic acid-binding proteins with nucleic acids, and the Y61D and M64D substitutions are known to disrupt DNA-dependent oligomerization of H-NS. The purified H-NS$_{Y61DM64D}$ protein was snap-frozen in liquid nitrogen and stored at -80 °C until further use.

**Data Analysis**

All imaging data were acquired and recorded using NIS Elements software and subsequently exported as TIFF files for further analysis. Subsequent analysis was performed using a custom-developed MATLAB (MathWorks) software pipeline. The Cy5 and SYTOX Orange (SxO) images were collected sequentially, acquiring 100 frames for the Cy5 channel and 1000 frames for the SxO channel. Both channels were co-registered to spatially correlate the Cy5 fluorescence with the SxO-stained DNA, with the Cy5 label positioned at one end of the DNA molecule serving as a fiducial marker to establish molecular directionality. At each position along the DNA contour, the local fluorescence intensity was computed by integrating pixel values across 11 adjacent pixels oriented perpendicular to the DNA axis, with background subtraction performed using the median intensity of the pixels flanking the molecule. The background-corrected intensity profiles were subsequently assembled into two-dimensional intensity kymographs by stacking successive intensity profiles along the time axis, enabling visualization and quantification of DNA and protein dynamics. These kymographs were used to monitor and analyze plectoneme formation and dynamics, nucleoprotein filament assembly along the DNA, and H-NS-mediated DNA bridging. Nucleoprotein filament formation was identified by a localized reduction in fluorescence intensity along the DNA, while DNA bridging by H-NS manifested as a localized fluorescence intensity enhancement, reflecting increased local DNA density at bridging sites. Plectoneme identification and spatial density quantification were performed following the procedures described previously[2,4]. Briefly, a threshold-based detection algorithm was applied to the DNA density profile. The median density computed across the full kymograph was used as the background reference, and a detection threshold of 25% above this baseline was applied. Only intensity peaks that persisted for a minimum of two consecutive time frames (200 ms) were designated as plectonemes, and the resulting plectoneme density profiles were computed and normalized. Plectoneme identification and spatial density quantification were performed following the analytical approach of, in which a threshold-based detection algorithm was applied to the DNA density profile. The median density computed across the full kymograph was used as the background reference, and a dynamic detection threshold above this baseline was applied. Only intensity peaks that persisted for a minimum of three consecutive time frames (≥300 ms) were designated as plectonemes, and the resulting plectoneme density profiles were computed and normalized.

**Plectoneme Occupancy**

Plectoneme occupancy was calculated as the fractional time a given genomic position was occupied by a plectoneme over the course of the experiment. To quantify this, a binary matrix was constructed in which each row corresponds to a genomic position, and each column corresponds to an individual time frame. Each element of the matrix was assigned a value of 1 if the position was covered by a plectoneme in that frame, or 0 otherwise. The plectoneme occupancy at each genomic position was then defined as:

$$\frac{1}{N}\sum_{t=1}^{N} b_{i,t}$$

Where $b_{i,t}$ is the binary value at genomic position $i$ at time frame $t$, and N is the total number of frames analyzed. This yields a value between 0 and 1, representing the fraction of time each position is occupied by a plectoneme.

**Plectoneme Diffusion**

To quantify the lateral mobility of DNA plectonemes, the time-averaged mean squared displacement (MSD) was calculated from plectoneme trajectories. Only trajectories meeting a minimum duration of 10 frames (1.0 s) and a mean plectoneme size exceeding 300 bp (0.3 kb) were included in the analysis to ensure statistical reliability.

The MSD for each trajectory was computed as:

$$\mathrm{MSD}(\tau) = \frac{1}{N-\tau}\sum_{t=1}^{N-\tau}[x(t+\tau) - x(t)]^2$$

To account for the inherent statistical correlation and the decrease in independent data points at larger time lags, we applied a non-linear least-squares fitting weighted by the reciprocal of the theoretical variance of the MSD, defined as

$$w(\tau) = \left[\frac{2\tau(N+1-\tau)}{3}\right]^{-1}$$

The diffusion coefficient was determined for each trajectory by fitting a linear model ($\mathrm{MSD}(\tau)=\alpha\tau$). To further characterize the movement, ensemble-averaged MSD was fit to both a linear model and a power law model ($\mathrm{MSD}(\tau) = \alpha\tau^n$) to account for sub-diffusive behavior.

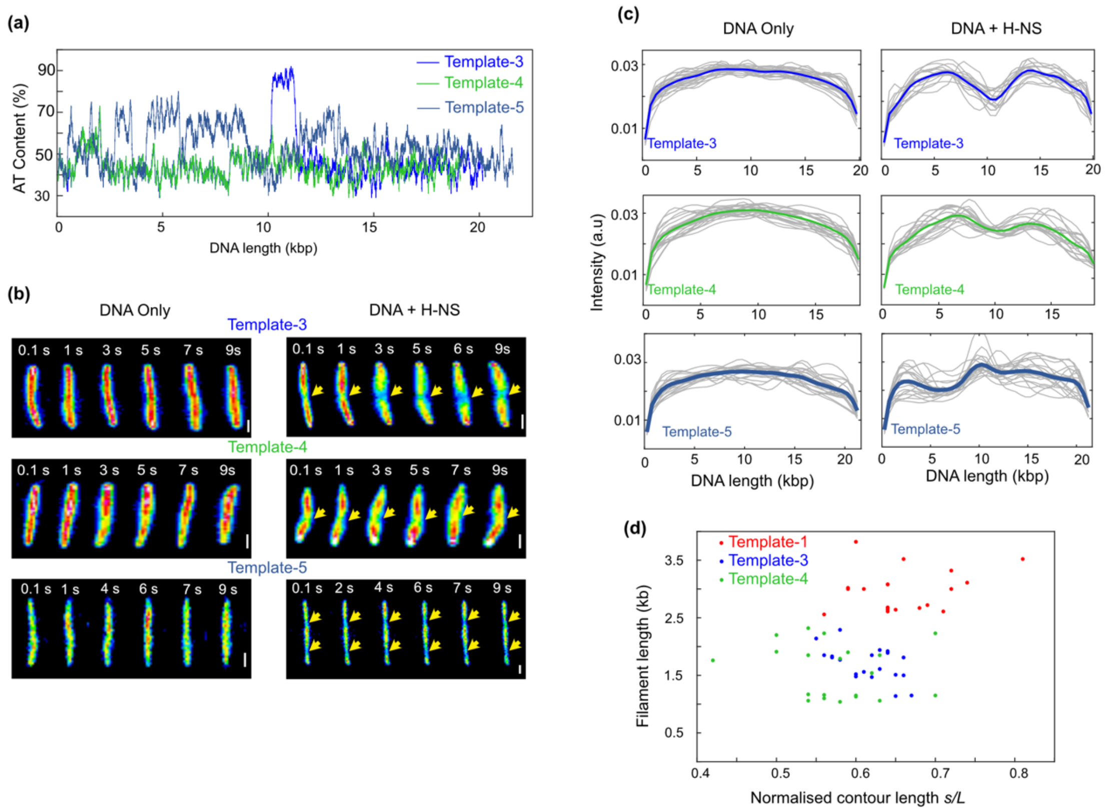


**Supplementary Figure-1: H-NS nucleoprotein filament formation on various DNA constructs.** **(a)** Graph displaying the AT percentage along the DNA length for Template-3: 1 kb 90% AT-rich segment at the center, Template-4: 100 bp H-NS nucleation site at the center and Template 5: Multi-AT-rich-loci (Blue, Green and Dark blue respectively), AT-content averaging window size is 100 bp. **(b)** Representative snapshots of relaxed DNA without (left) and with H-NS (right) for Template - 3, -4 and -5. The fluorescence depleted regions in the DNA+H-NS column indicate filament formation at AT-rich loci which are marked with yellow arrows. **(c)** Average fluorescence intensity profiles along normalized DNA contour without (left) and with H-NS (right) for Template- 3, 4 and 5. Grey: individual molecules, and Blue, green, and dark blue mean for Template-3, 4, and 5 respectively, each frame- 100 msec. **(d)** Scatter plot displaying length of the H-NS nucleoprotein filament on Template 1, 3 and 4 along the normalized length (end-to-end-length/contour length). Each dot represents one molecule. n ≥ 20 molecules for each data (c and d); Scale Bar: 1 µm (b).

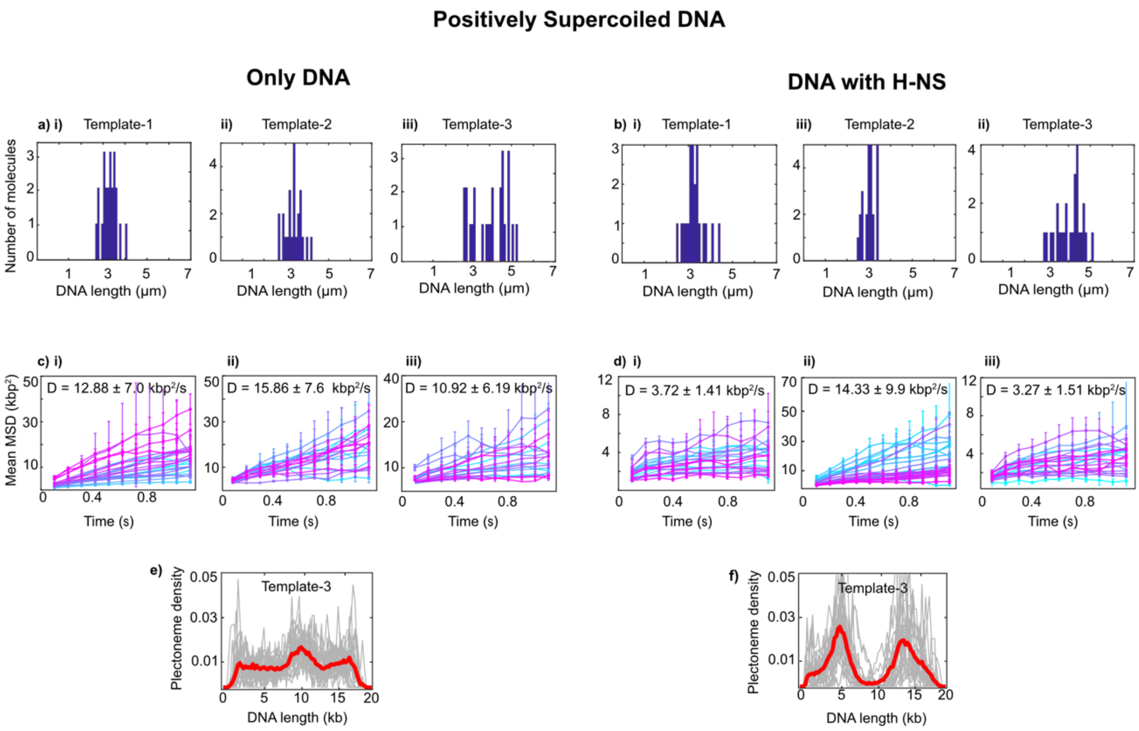


**Supplementary Figure-2: Behavior of H-NS on positively supercoiled DNA on various DNA constructs. (a) i, ii,** and **iii** -End-to-end length distribution of individual DNA molecules without H-NS for Template-1, -2 and -3 respectively, where the x-axis represents the end-to-end length of each DNA molecule and the y-axis represents the number of molecules **(b) i, ii, and iii-** End-to-end length distribution of individual DNA molecules for Template- 1, 2 and 3 respectively with H-NS **(c) i, ii, iii-** Mean square displacement (MSD) over time for Template- 1, 2 and 3, respectively, without H-NS. MSD for a number of plectonemes in a molecule is averaged and plotted against time in seconds each color represents each molecule, D-represents average diffusion coefficient of all the molecule. **(d) i, ii, and iii-** Mean square displacement (MSD) over time for Template- 1, 2 and 3 respectively with H-NS. **(e)** Plectoneme density over 1000 frames along the DNA length for Template-3 without H-NS and **(f)** with H-NS. n ≥ 20 molecules for all plots; grey: individual molecules and red: mean.

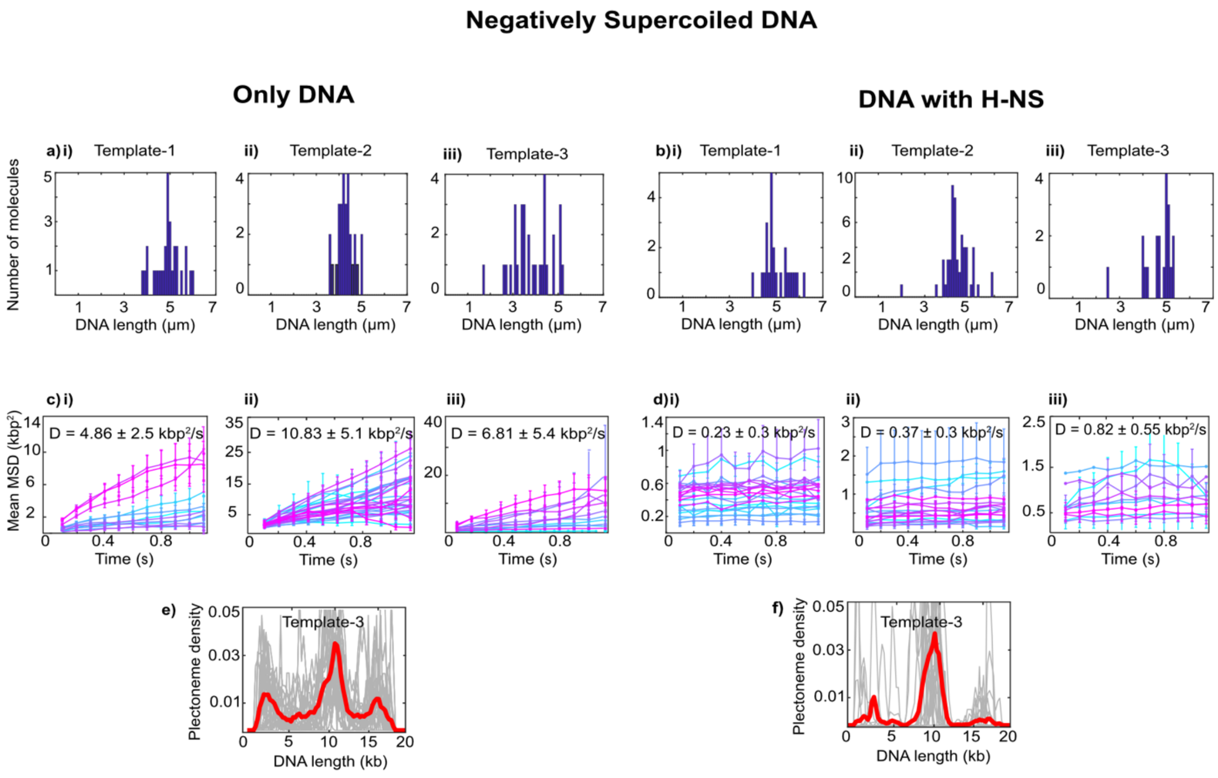


**Supplementary Figure-3: Behavior of H-NS on negatively supercoiled DNA on various DNA constructs. (a-) i, ii, and iii-** End-to-end length distribution of individual DNA molecules without H-NS for Template- 1, 2 and 3 respectively, where the x-axis represents the end-to-end length of each DNA molecule and the y-axis represents the number of molecules **(b) i, ii, and iii-** End-to-end length distribution of individual DNA molecules for Template- 1, 2 and 3 respectively with H-NS **(c) i, ii, and iii-** Mean square displacement (MSD) over time for Template- 1, 2 and 3 respectively without H-NS. MSD for a number of plectonemes in a molecule is averaged and plotted against time in seconds each colour represents each molecule, D-represents average diffusion coefficient of all the molecule. **(d) i, ii, and iii-** Mean square displacement (MSD) over time for Template- 1, 2 and 3 respectively with H-NS. **(e)** Plectoneme density over 1000 frames along the DNA length for Template-3 without H-NS and **(f)** with H-NS. n ≥ 20 molecules for all plots; grey: individual molecules and red: mean.

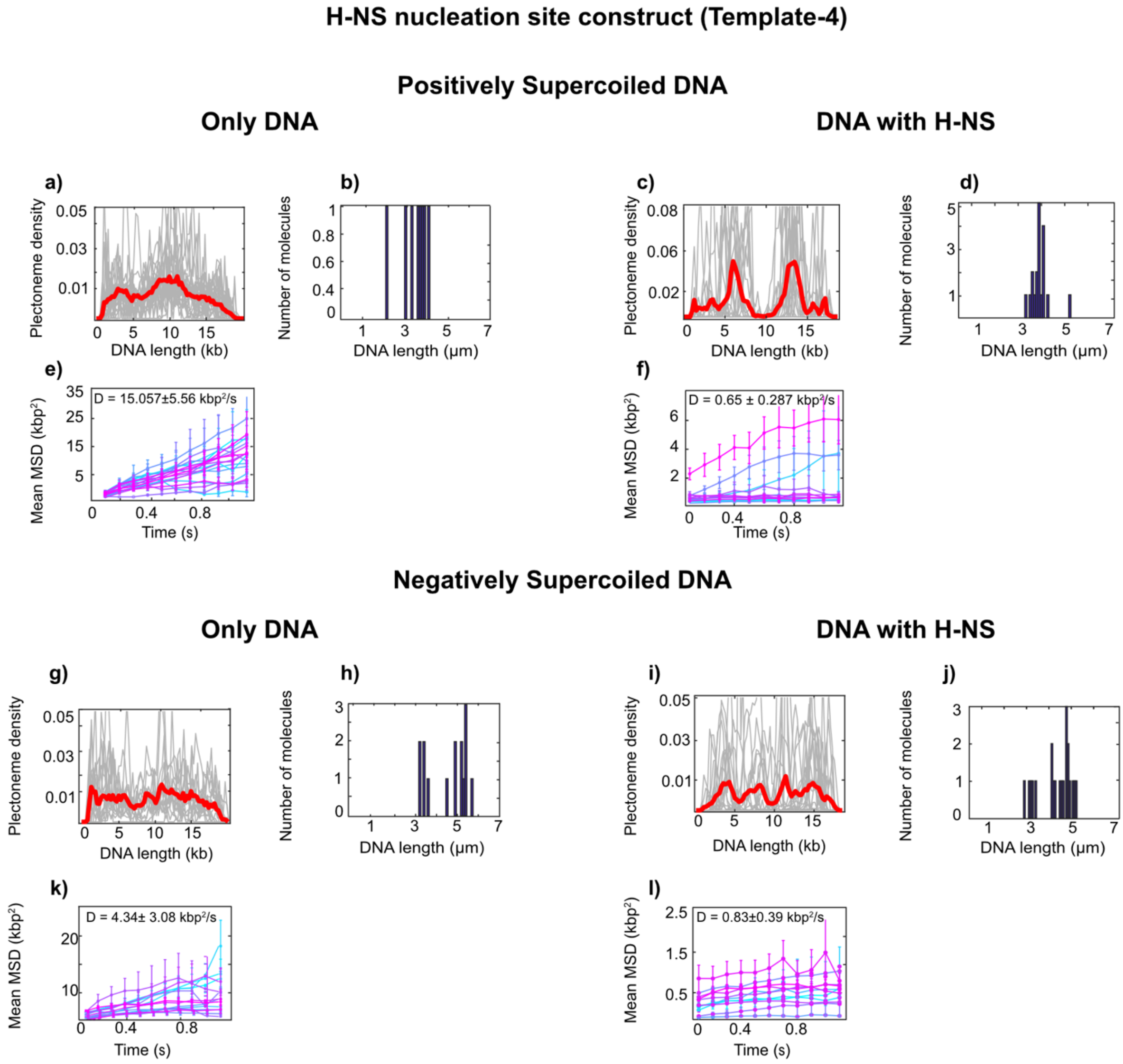


**Supplementary Figure-4: Behavior of H-NS on positively and negatively supercoiled DNA on a DNA construct with 100 bp H-NS nucleation site at the center. (a-f)** Quantitative analysis of H-NS behavior on positively supercoiled DNA **(a)** Plectoneme density over 1000 frames along the DNA length without H-NS. **(b)** End-to-end length distribution of individual DNA molecules without H-NS. **(c)** Plectoneme density over 1000 frames along the DNA length with H-NS. **(d)** End-to-end length distribution of individual DNA molecules with H-NS. **(e-f)** Mean square displacement (MSD) over time without and with H-NS respectively. **(g-l)** Quantitative analysis of H-NS behavior on negatively supercoiled DNA. **(g)** Plectoneme density over 1000 frames along the DNA length without H-NS. **(h)** End-to-end length distribution of individual DNA molecules without H-NS, where the x-axis represents the end-to-end length of each DNA molecule and the y-axis represents the number of molecules. **(i)** Plectoneme density over 1000 frames along the DNA length with H-NS. $n \geq 20$ molecules for all data; grey: individual molecules, red: mean. **(j)** End-to-end length distribution of individual DNA molecules with H-NS. **(k)** Mean square displacement (MSD) over time without and with H-NS respectively.

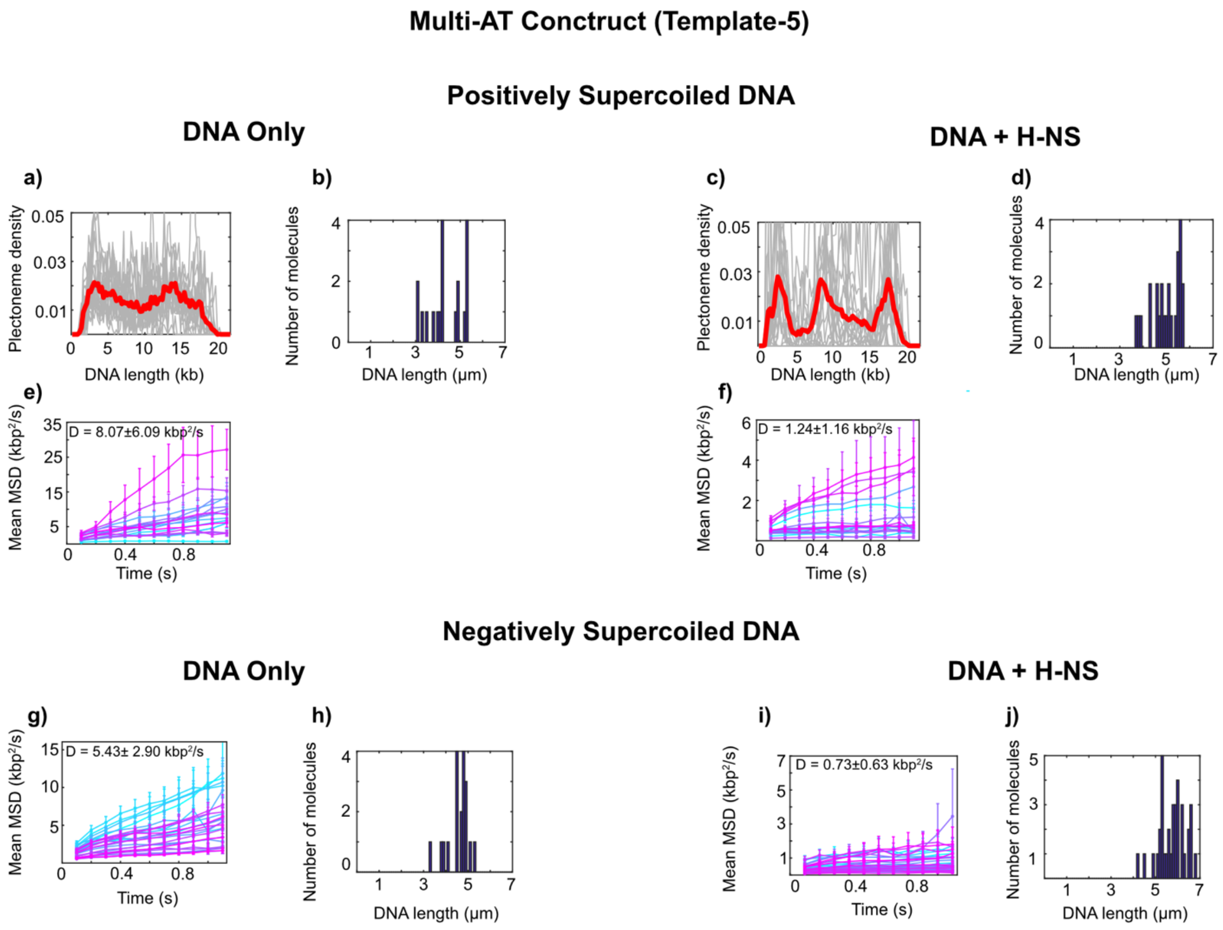


**Supplementary Figure-5: Behavior of H-NS on positively and negatively supercoiled DNA on a DNA construct with multiple AT rich sites (a-f)** Quantitative analysis of H-NS behaviour on positively supercoiled DNA **(a)** Probability of plectoneme formation over 1000 frames along the DNA length without H-NS. **(b)** End-to-end length distribution of individual DNA molecules without H-NS, where the x-axis represents the end-to-end length of each DNA molecule and the y-axis represents the number of molecules. **(c)** Probability of plectoneme formation over 1000 frames along the DNA length with H-NS. **(d)** End-to-end length distribution of individual DNA molecules with H-NS. **(e-f)** Mean square displacement (MSD) over time without and with H-NS respectively. **(g-j)** Quantitative analysis of H-NS behavior on negatively supercoiled DNA **(g)** Mean square displacement (MSD) over time without H-NS. **(h)** End-to-end length distribution of individual DNA molecules without H-NS, where the x-axis represents the end-to-end length of each DNA molecule and the y-axis represents the number of molecules. **(i)** Mean square displacement (MSD) over time with H-NS. **(j)** End-to-end length distribution of individual DNA molecules with H-NS.

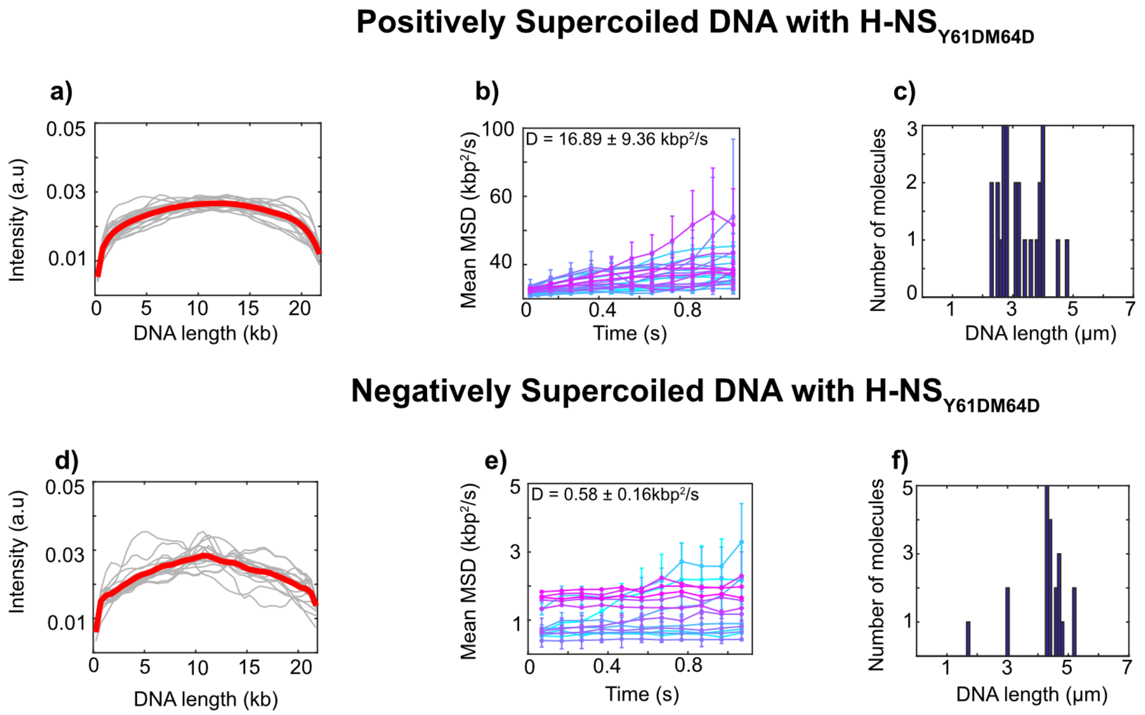


**Supplementary Figure-6. The oligomerization-deficient H-NS$_{Y61DM64D}$ mutant retains bridging mode of binding but cannot form stiffening filaments. (a-c)** Quantitative analysis of H-NS$_{Y61DM64D}$ behavior on positively supercoiled DNA **(a)** Average fluorescence intensity along the DNA length over 1000 frames. **b)** Mean square displacement (MSD) over time. **(c)** End-to-end length distribution of individual DNA molecules, where the x-axis represents the end-to-end length of each DNA molecule and the y-axis represents the number of molecules. **(d-f)** Quantitative analysis of H-NS$_{Y61DM64D}$ behavior on negatively supercoiled DNA **(d)** Average fluorescence intensity along the DNA length over 1000 frames. (**e)** Mean square displacement (MSD) over time. **(f)** End-to-end length distribution of individual DNA molecules. n ≥ 20 molecules; grey: individual molecules, red: mean.

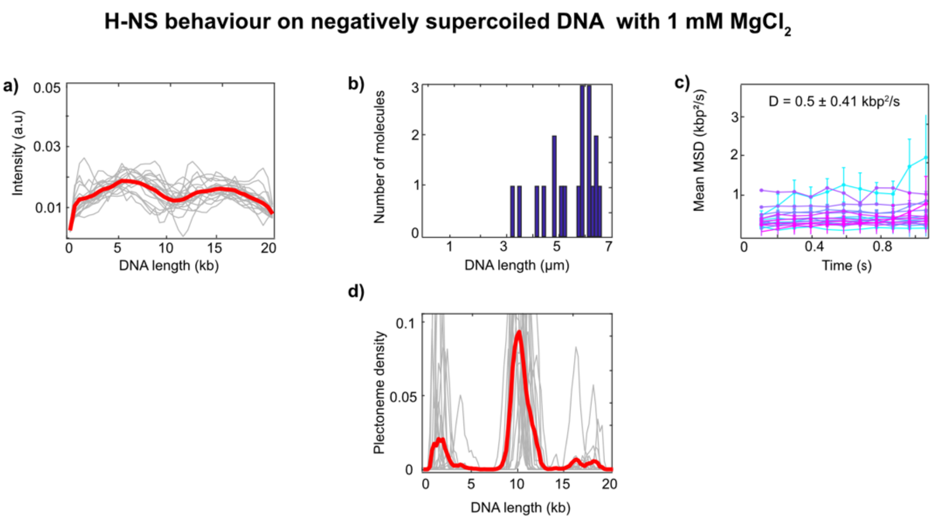


**Supplementary Figure 7: H-NS behavior on negatively supercoiled Template-1 DNA with 1 mM $MgCl_2$. (a)** Average fluorescence intensity along the DNA length over 1000 frames. **(b)** End-to-end length distribution of individual DNA molecules, where the x-axis represents the end-to-end length of each DNA molecule and the y-axis represents the number of molecules. **(c)** Mean square displacement (MSD) over time. (**d)** Probability of plectoneme formation over 1000 frames along the DNA length with H-NS. $n \geq 20$ molecules; grey: individual molecules and red: mean.

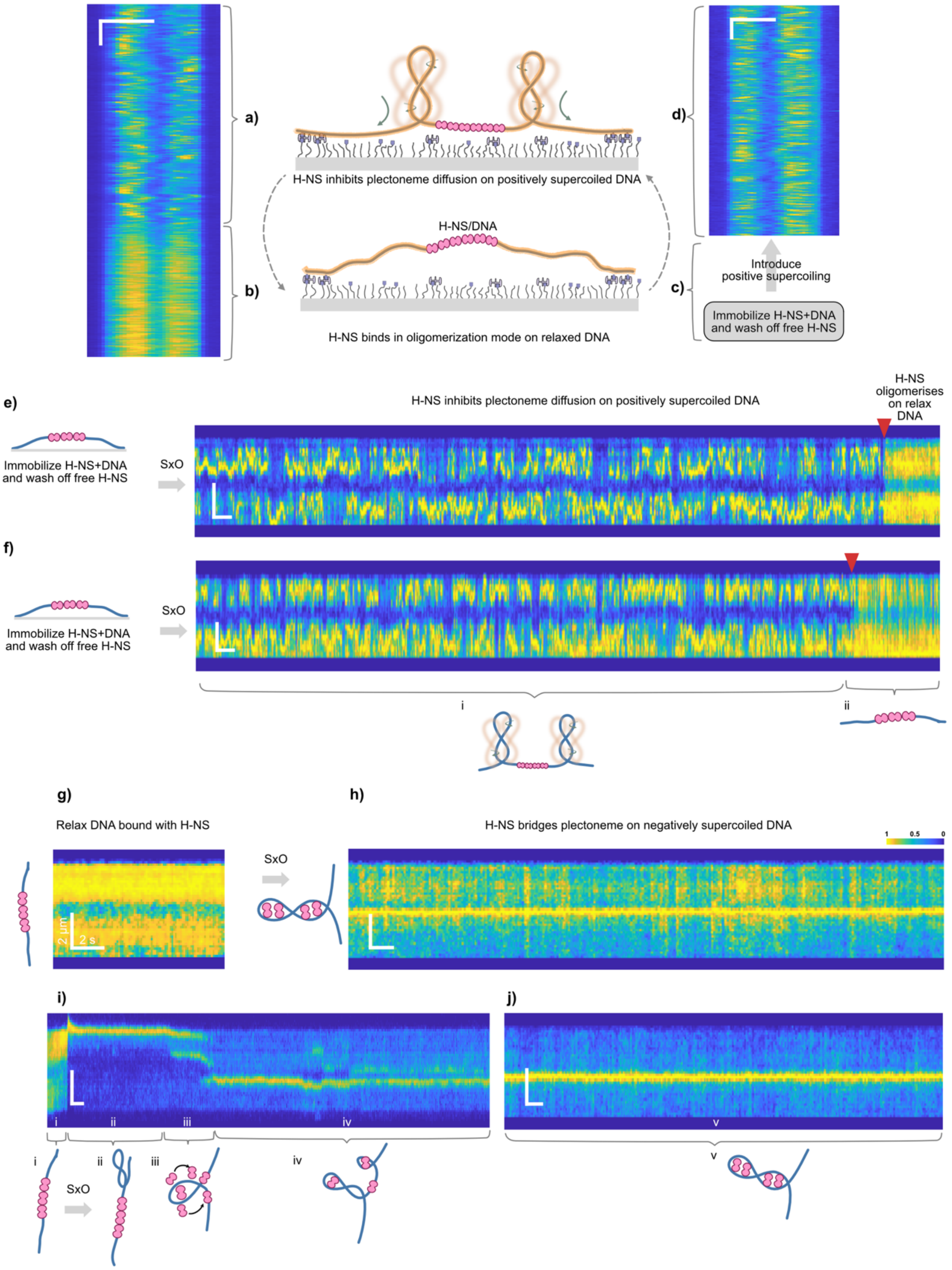


**Supplementary Figure 8. H-NS switches binding mode upon change in DNA helicity**. **(a-d)** Transition from relax to positively supercoiled DNA and vice versa with bound H-NS on Template-3 maintains the filament mode and acts as topological barrier **(a)** H-NS nucleoprotein filament acts as a topological barrier on positively supercoiled DNA with dark region at the centre representing H-NS nucleoprotein filament.**(b)** upon photoinduced nicking the DNA relaxes thereby no diffusion of plectonemes are observed while nucleoprotein filament is intact on the DNA. **(c)** The DNA and H-NS was incubated in the tube for 5 minutes and flowed into the channel. Excess H-NS was washed off from the channel. **(d)** SxO was flowed into the channel to generate positively supercoiled DNA. The H-NS nucleoprotein filament appears as dark region at AT-rich site while the plectonemes are diffusing along the AT-poor flanking sites. **(e)** and **(f)** represent examples for positively supercoiled DNA to relaxed DNA with bound H-NS. Upon photoinduced nicking (red arrow) the DNA relaxes thereby no diffusion of plectonemes are observed while nucleoprotein filament is intact on the DNA. **(g–h)** Stiffening-to-bridging switch triggered by introducing negative supercoiling via SxO dilution. **(g)** Stiffening filament (fluorescence depleted region in the center) on relaxed Template-1 at high SxO. **(h)** SxO dilution induces negative supercoiling; fluorescence depleted region is replaced by a bridging punctum. **(i-j)** Real-time transition event from stiffening to bridging mode. **(i)-i** Stiffening filament (fluorescence depleted region) on relaxed DNA at high SxO concentration. **ii** Low SxO concentration introduced into the channel leading to negative supercoiling of DNA. The plectoneme is pushed towards one side with the fluorescence depleted region at the center. **iii** and **iv** H-NS transitions into bridging mode and stabilize plectoneme at the center. **(j)** Plectoneme is bridged at the center at the AT-rich site. Horizontal scale Bar: 2 μm and vertical scale bar: 2 s (figure: a-c). Vertical scale Bar: 2 μm and horizontal scale bar: 2 s (figure: e-j)

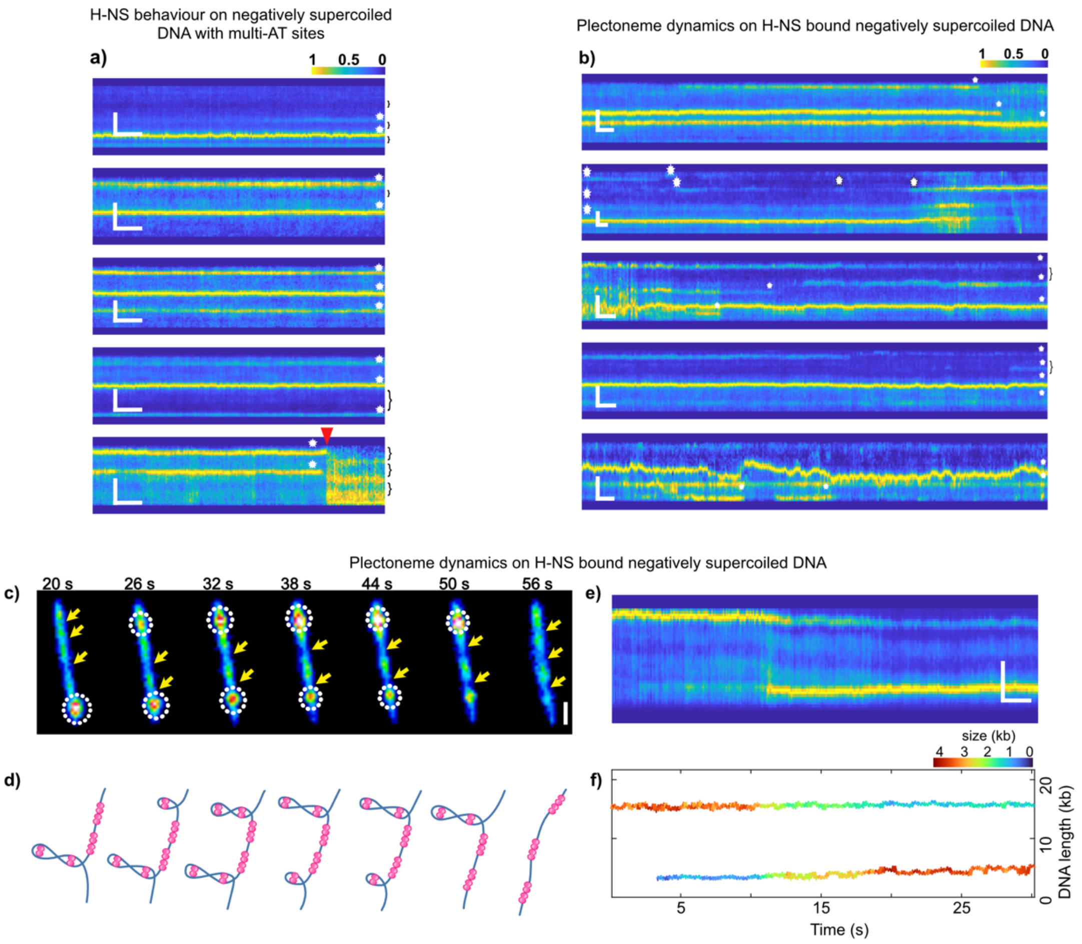


**Supplementary Figure-9. Helicity-dependent H-NS organization drives locus-specific silencing on multi-AT site DNA. (a)** Representative fluorescence kymographs of Template-5 under negatively supercoiled in the presence of H-NS. White dots (puncta) indicate H-NS bridging events of plectoneme, while brackets denote regions where H-NS binds in the filamentous mode, leading to fluorescence depletion. Red arrowhead marks the point at which the DNA transitions to a relaxed state, whereupon H-NS reorganizes into the stiffening binding mode, confirming that H-NS binding mode at multi-AT loci is strictly dependent on DNA super helicity. **(b-f)** Dynamics of plectonemes on a molecule for multi-AT locus sites DNA. **(b)** Representative fluorescence kymographs of Template-5 in the presence of H-NS showing dynamic rearrangement of bridged plectonemes. White dots (puncta) indicate H-NS bridging events, and brackets denote regions of filamentous H-NS binding, illustrating stochastic switching between these two modes at discrete AT-rich loci. **(c)** Representative snapshots for plectoneme dynamics on Template-5 in which an H-NS bridged plectoneme gradually dissipates at one position while growing at another position. White circles represent plectonemes and yellow arrows represent fluorescence depleted regions due to H-NS oligomerization. **(d)** Schematic depiction for the molecule in figure (c). **(e)** Another representative kymograph for over 300 frames showing plectoneme dynamics along the DNA contour. **(f)** Plectoneme size distribution along each frame corresponding to figure (e). Vertical scale Bar: 2 µm (a-d) and horizontal scale bar: 2 s (a-d)

**Supplementary Figure-10**

HNS Oligomerization-deficient mutant (Y61DM64D) on Multi-AT Construct Negatively Supercoiled DNA.

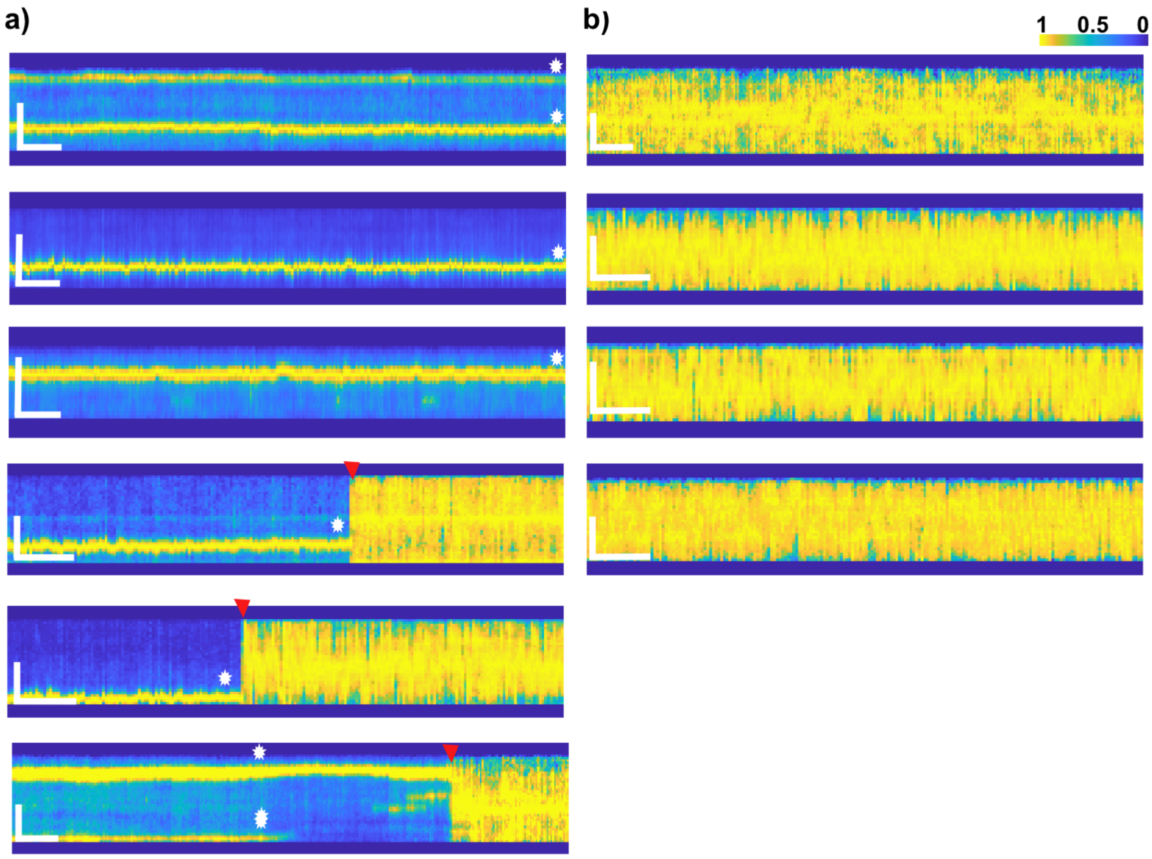


**Supplementary Figure-10. Oligomerization-deficient H-NS$_{Y61DM64D}$ retains bridging activity but fails to form filaments on Template-5 DNA. (a)** Representative fluorescence kymographs of Template-5 under negatively supercoiled conditions in the presence of oligomerization-deficient mutant H-NS$_{Y61DM64D}$. White dots (puncta) indicate H-NS bridging events at AT-rich loci, confirming that the mutant retains bridging activity. In the bottom three kymographs, red arrow heads mark the point at which the DNA transitions to a relaxed state, whereupon the absence of any local reduction in fluorescence intensity clearly indicates that H-NS$_{Y61DM64D}$ fails to bind in the filamentous mode, confirming that filament formation is strictly dependent on oligomerization capacity. **(b)** Representative fluorescence kymographs of relaxed Template-5 DNA in the presence of H-NS$_{Y61DM64D}$. The homogeneous fluorescence intensity distribution along the DNA contour confirms that H-NS$_{Y61DM64D}$ is unable to form filaments, consistent with its oligomerization-deficient nature.

### Supplementary movie legends

**Movie S1.** Movie showing SxO stained double tethered relaxed DNA for Template-1. SxO is homogenously distributed providing a uniform fluorescence intensity along the length of the DNA (Template-1) in the absence of H-NS.

**Movie S2.** Movie showing positively supercoiled DNA for Template-1. The plectoneme appear as high intensity punctum which diffuses all along the DNA contour in the absences of H-NS.

**Movie S3.** Movie showing positively supercoiled DNA for Template-1 with H-NS. The nucleoprotein filament at the centre acts as a topological barrier for the diffusion of plectonemes
**Movie S4.** Movie showing negatively supercoiled DNA for Template-1 bound with H-NS, plectoneme stabilised at the AT rich site appearing as high intensity punctum at the centre.
**Movie S5.** Movie showing SxO stained double tethered relaxed DNA for Template-2. SxO is homogenously distributed providing a uniform fluorescence intensity along the length of the DNA in the absence of H-NS.

**Movie S6.** Movie showing positively supercoiled DNA for Template-2. The plectoneme appear as high intensity punctum which diffuses all along the DNA contour in the absences of H-NS.

**Movie S7.** Movie showing positively supercoiled DNA (Template-2) bound with H-NS, no dark region at the AT rich site. Plectoneme diffuses freely along the DNA contour.

**Movie S8.** Movie showing negatively supercoiled DNA (Template-2) bound with H-NS, plectoneme stabilised at multiple sites appearing as high intensity punctum.

**Movie S9.** Movie showing negatively supercoiled DNA (Template-1) bound with H-NS, plectoneme stabilised at the AT rich site appearing as high intensity punctum at the centre. During this imaging, flow was introduced to show the high intensity punctum at the centre is a plectoneme which is stretched under the laminar flow

***Supplementary Table 1: Primers list***

| Construct Name | Primer Name | Sequence |
|---|---|---|
| Ec H-NS | Ec H-NS_FP | CTTTAAGAAGGAGATATACCATGAGCGAAGCACTTAAAATTC |
| | Ec H-NS_RP | CAGTGGTGGTGGTGGTGGTGTTGCTTGATCAGGAAATCGTC |
| Ec H-NS Y61D | Ec H-NS Y61D_FP | ACTGCAGCAAGATCGCGAAATGCTGATCGCT |
| | Ec H-NS Y61D_RP | GCATTTCGCGATCTTGCTGCAGTTTACGAGT |
| Ec H-NS M64D | Ec H-NS M64D_FP | ATATCGCGAAGATCTGATCGCTGACGGTATT |
| | Ec H-NS M64D_RP | CAGCGATCAGATCTTCGCGATATTGCTGCAG |
| Template 1 | 3 kb AT+XbaI_FP | TTTTTTTTTTCTAGAGCAGAGCTGGAAGTGCAGAC |
| | 3 kb AT+PciI_RP | TTTTTTTTTTACATGTAAACATCCCTTACACTGGTG |
| Template 2 | 3kb_moderate AT_XbaI_18kb DNA F.P | TTTTTTTTTTCTAGAGCAGAGCTGGAAGTGCAGAC |
| | 3Kb_moderate AT_PciI_18Kb R.P | TTTTTTTTTTACATGTAAACATCCCTTACACTGGTG |
| Template 4 | HNS_100bp_XbaI_FP | TTTTTTTTTTCTAGAGTTCCCGGTACCCTGGTC |
| | HNS_100bp_HindIII_RP | TTTTTTTTTTAAGCTTTCTTCAACGCTGTATGTATAG |
| 500 bp Biotin Handles | BiotinH_FP | GGTTTGCGTATTGGGCGCTCTTCCG |

| | | |
|---|---|---|
| | BiotinH_RP_NotI | GCGGCCGCAGACGATAGTTA CCGGATAAGGCGC |
| | BiotinH_RP_XhoI | CTCGAGAGACGATAGTTACCG GATAAGGCGC |


1 Gaur, P. *et al*. Sequence-dependent co-condensation of Lsr2 with DNA elucidates the mechanism of genome compaction in Mycobacterium tuberculosis. *Nucleic Acids Research* **54** (2026). https://doi.org/10.1093/nar/gkaf1428

2 Kim, S. H. *et al.* DNA sequence encodes the position of DNA supercoils. *Elife* **7** (2018). https://doi.org/10.7554/eLife.36557

3 Chandradoss, S. D. *et al*. Surface passivation for single-molecule protein studies. *J Vis Exp* (2014). https://doi.org/10.3791/50549

4 Ganji, M., Kim, S. H., van der Torre, J., Abbondanzieri, E. & Dekker, C. Intercalation-Based Single-Molecule Fluorescence Assay To Study DNA Supercoil Dynamics. *Nano Letters* **16**, 4699–4707 (2016). https://doi.org/10.1021/acs.nanolett.6b02213
